# Dynamic Tumor Immune Microenvironment Remodeling Predicts Response of Checkpoint Inhibitor Therapy

**DOI:** 10.1101/2024.06.30.601413

**Authors:** Zhongyang Lin, Mitalee Chandra, Nalini Srinivas, Selma Ugurel, Jürgen C. Becker, Dvir Aran

## Abstract

Immune checkpoint inhibitors (ICI) have transformed cancer therapy, yet the basis of variable patient responses remains unclear. We assembled a longitudinal single-cell RNA sequencing atlas of 441 samples from 241 patients across ten cancers to map treatment-associated remodeling of the tumor immune microenvironment (TIME). Using a hierarchical reference-guided deep-phenotyping framework, we defined 77 immune and stromal subtypes and resolved four conserved TIME subtypes. Approximately 40% of tumors shifted between states during therapy, and the transition was more predictive of outcome than the baseline state. Across 1,988 bulk transcriptomic tumors, favorable transitions toward inflamed or B-cell-enriched subtype tracked with improved response and survival, while persistence in or shifts towards myeloid dominance indicated resistance. We derived a transition score that predicted outcomes for baseline tumors across independent cohorts. These findings establish immunotype transitions as a central determinant of ICI, offering new avenues for response prediction and rational immunotherapy design.

**Highlights:**

- A pan-cancer meta-analysis maps treatment-associated remodeling of the tumor immune microenvironment.
- Four conserved TIME states emerge across cancer subtypes.
- TIME transition patterns during treatment are associated with clinical outcome.
- Baseline immune programs derived transition scores predict treatment response and survival.

## Introduction

Immune checkpoint inhibitors (ICIs) transformed cancer therapy by reinvigoration of anti-tumor immunity, achieving durable responses in patients with previously intractable malignancies.^1–5^ Thus, ICIs have become standard-of-care for many cancer types. Despite the successes, however, more than half of the patients either fail to respond initially or develop acquired resistance. This notion has driven an intense search for biomarkers to improve response prediction and allow personalized therapeutic strategies.

The tumor immune microenvironment (TIME) critically impacts the outcome of anti-tumor immune responses.^6^ Tumors are often described along a hot-to-cold spectrum that reflects the distribution and functional state of immune cells within and around the tumor. Hot tumors typically show cytotoxic T cells, interferon signaling, and antigen presentation features, and tend to respond better.^7^ Cold tumors tend to be T-cell-poor or T-cell-excluded and are often dominated by suppressive myeloid and regulatory programs that resist activation.^8^ These patterns have been observed with bulk expression, spatial assays, and single-cell profiling and have anchored many current predictors.^9–13^ Yet this framework implicitly treats the TIME as a fixed, pre-treatment attribute of the tumor. Checkpoint blockade, however, is not a passive test of immune readiness but a potent perturbation that can rewire cellular composition and function. This simple assumption might overlook the fact that ICI acts as a profound perturbation, actively reshaping the TIME by reinvigorating lymphocytes and altering myeloid populations. Early on-treatment molecular remodeling, including T-cell reinvigoration, B-cell activation, and clonal expansion, has been shown to correlate with clinical benefit.^14–17^ Recognizing the TIME as a dynamic system rather than a static trait reframes the question from whether a tumor is hot or cold to how its immune ecosystem evolves once therapy begins and whether those trajectories can predict outcome.^18^

Testing this hypothesis requires longitudinal sampling of tumors before and during therapy, enabling direct observation of TIME remodeling and its link to treatment efficacy. Analyses on consecutively sampled tumors have provided unprecedented insights into TIME dynamics in easy-to-biopsy cancers;^15,16,19^ this approach has been expanded to include the neoadjuvant therapy setting, which allows for the collection of sequential tumor samples.^20–30^ Nonetheless, investigating TIME dynamics has faced several challenges. For example, studies examining treatment-associated compositional shifts typically include 10 to 20 patients with paired samples, lacking sufficient statistical power to detect subtle but clinically relevant features. Comparing data across different studies is hampered by batch effects and methodology inconsistencies, such as cell annotation and compositional analysis. Thus, although the TIME cell lineages and classifications have been extensively explored,^9,10,31–38^ the full plot of TIME dynamics and the association with treatment efficacy remain poorly characterized. Still, pan-cancer meta-analysis holds great potential once the challenge of data harmonization has been solved.

Here, we address this gap by assembling the largest longitudinal single-cell RNA sequencing compendium of ICI-treated tumors to date, encompassing 441 samples from 241 patients across ten cancer types. To overcome the challenges of cross-cohort integration, we developed a hierarchical, reference-guided deep-phenotyping framework that enables consistent cell-type classification and compositional analysis across studies. This approach allowed us to define a compact and reproducible set of TIME states and to track transitions between them in paired pre-and on-treatment samples. We hypothesized that the direction of TIME remodeling during ICI therapy, rather than the baseline state alone, dictates clinical outcome, and that baseline molecular programs determine each tumor’s propensity for favorable or unfavorable evolution. By mapping these dynamics across cancer types, we reveal conserved trajectories of immune remodeling that link early cellular plasticity to therapeutic efficacy.

## Results

### Pan-cancer atlas reveals landscape of tumor immune microenvironment dynamics

To systematically investigate cellular remodeling under ICI therapy, we first generated scRNA-seq data from paired tumor biopsies of 17 melanoma patients collected at treatment initiation (day 0) and 6-16 days after anti-PD-1 ± anti-CTLA-4 therapy initiation, capturing early treatment-associated changes (**Table S1**). We further assembled a pan-cancer atlas by integrating sequential scRNA-seq datasets from 15 independent studies encompassing ten cancer types. A key feature of this atlas is the inclusion of consecutively sampled, site-matched tumors from ICI-treated patients, covering a diverse range of malignancies: cutaneous melanoma (SKCM), head and neck squamous cell carcinoma (HNSCC), non-small cell lung cancer (NSCLC), ER+ breast cancer (BRCA), triple-negative breast cancer (TNBC), hepatocellular carcinoma (HCC), colorectal cancer (CRC), prostate cancer (PC), and basal cell carcinoma (BCC) (**Figure 1A-C, Table S2**). After rigorous quality control, the atlas comprised 1,483,740 TIME cells derived from 370 samples with complete profiling and 71 samples from CD45+ sorted cells.

**Figure 1.**
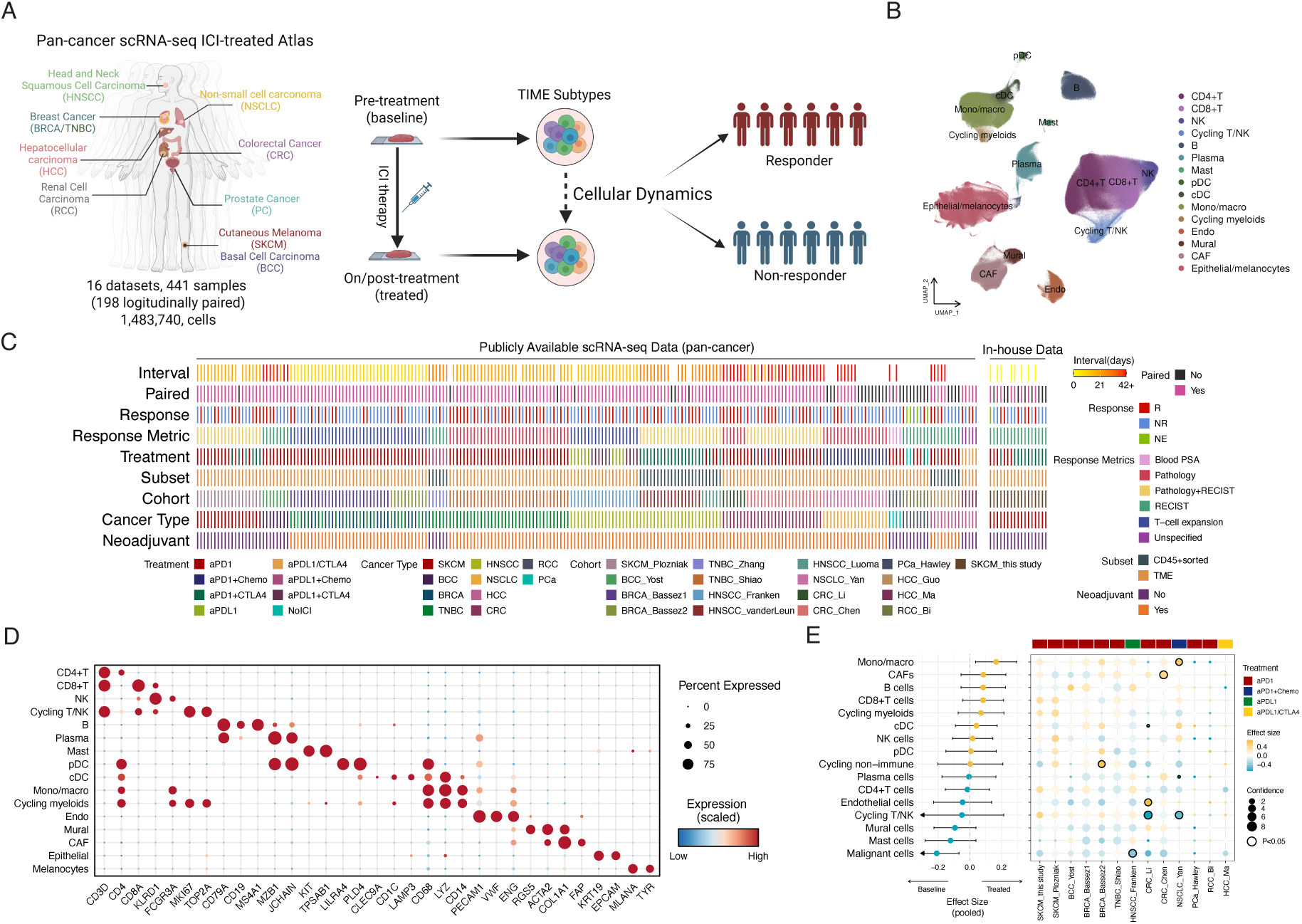
Construction and composition of the longitudinal single-cell atlas of ICI-treated tumors. **(A)** Schematic overview of the study design and analytical workflow, illustrating integration of longitudinal scRNA-seq datasets from multiple cancer types and mapping of tumor immune microenvironment (TIME) dynamics between pre- and on-treatment samples. **(B)** Uniform manifold approximation and projection (UMAP) of the integrated dataset, showing major immune and stromal lineages across all samples. **(C)** Summary of included studies and metadata, indicating cohort origin, cancer type, treatment regimen, sampling interval, and availability of paired and response information. **(D)** Validation of main cell-type assignments, with canonical marker gene expression confirming accurate classification of eleven distinct cell lineages. **(E)** Meta-analysis of compositional shifts during ICI therapy, showing pooled effect sizes for each major cell type (forest plot) and study-specific estimates (dot plots). Statistical significance was assessed using paired Wilcoxon signed-rank tests.

We applied a hierarchical reference-guided deep-phenotyping framework to assign cell identities at multiple resolutions. In the first step, each dataset was independently annotated with SingleR, supplemented by scGate filtering and Seurat clustering to resolve ambiguous populations. These preliminary labels were then iteratively refined throughout the annotation: misclassified clusters were corrected based on marker expression and potential doublets were removed (**Methods**). Finally, we performed cross-dataset integration within each major lineage to harmonize labels and ensure consistency across cohorts (**Figure 1B and Figure S1A-C**). Canonical marker gene expression confirmed accurate classification of 11 distinct cell types across all datasets (**Figure 1D and Figure S1D**).

To investigate treatment-associated changes in cell composition, we performed a cross-cohort meta-analysis of paired baseline and on-treatment samples (**Methods**). This analysis revealed a consistent enrichment of monocytes/macrophages paralleled by a reduction in malignant cells upon ICI therapy (**Figure 1E**). These compositional trends were reproducible across studies and correlated with standardized binary response metrics (**Figure S2A-B**). Beyond these global patterns, cancer-type-specific dynamics emerged. For instance, B cell infiltration increased in BCC, BRCA and HNSCC, whereas plasma cells were mainly accumulated in baseline, particularly in NSCLC patients receiving anti-PD1 therapy combined with chemotherapy (**Figure S2C**). Subgroup analyses further refined these observations: in ER+ BRCA, a single cycle of anti-PD1 therapy increased myeloid cell infiltration, whereas in TNBC it increased cycling T/NK cells and reduced mast cells (**Figure S2D**). When stratified by T cell expansion, patients with expansion displayed elevated plasmacytoid dendritic cells (pDC) and B cells, while those without expansion had increased monocytes/macrophages and fewer CD8+ T cells; mast cell depletion was also observed in responders by RECIST (**Figure S2E**). In HNSCC, combination therapy showed a more uniform pattern, with accumulation of B cells, plasma cells, and mast cells within two weeks, whereas after two cycles (approximately 3-4 weeks), there was a significant reduction in pDC and classic DCs (cDC) (**Figure S2F**). Although these early compositional shifts were not significantly linked to T-cell expansion or short-term clinical response, they indicate that ICI induces widespread, multi-lineage remodeling of the TME extending well beyond T-cell reactivation.

### High-resolution immune cell mapping uncovers treatment- and response-linked TIME shifts

In the second step of our deep-phenotyping framework, we applied lineage-specific reference panels derived from curated pan-cancer single-cell atlases to resolve fine-grained immune and stromal subtypes within each lineage (**Methods; Figure S1A and 2A**). This hierarchical approach allowed accurate discrimination of overlapping populations such as T, NK, and myeloid subsets, while controlling for technical and study-specific biases (**Figure 2B**). The annotation framework identified 62 immune and 15 non-immune cell subtypes (**Table S3**). The assignment was confirmed by both marker gene expression and functional enrichment analysis with curated signatures, consistent with prior studies (**Figure S3 and S4**).^33,37,39–42^

**Figure 2.**
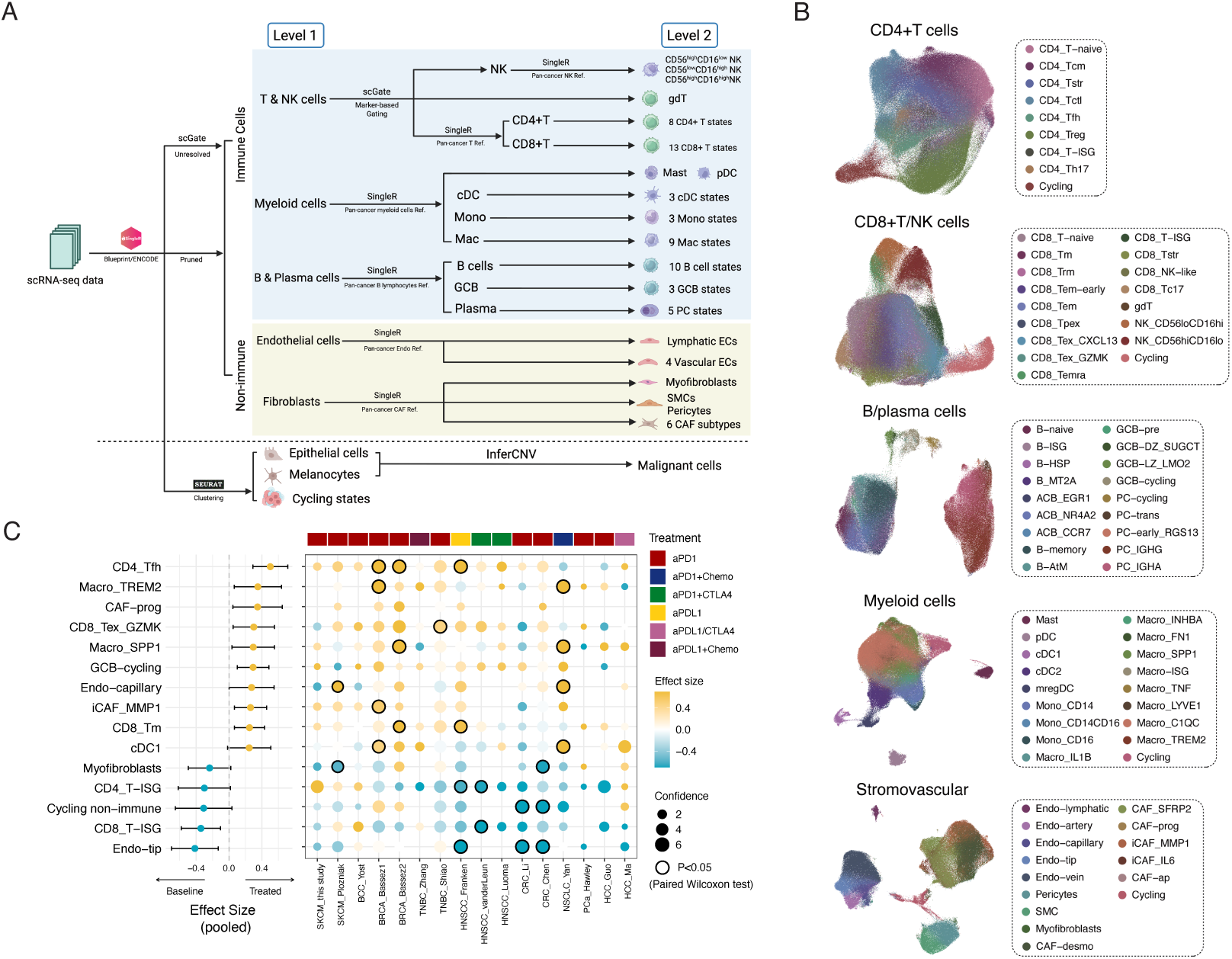
Hierarchical annotation framework and pan-cancer immune landscape. **(A)** Schematic overview of the two-level reference-guided deep-phenotyping workflow used for hierarchical cell annotation. **(B)** Uniform manifold approximation and projection (UMAP) visualization of fine-grained immune and stromal subtypes identified across the integrated pan-cancer atlas. **(C)** Meta-analysis of compositional changes during immune-checkpoint inhibitor (ICI) therapy. Forest plot shows the pooled effect size (rank-biserial correlation) for subtype abundance between baseline and treated tumors, with accompanying dot plots indicating study-specific estimates and confidence intervals. Statistical testing was performed using paired Wilcoxon signed-rank tests. Cell types abbreviations are available in **Table S3**.

We performed a meta-analysis of compositional changes across the atlas to identify consistent shifts during ICI therapy (**Figure 2C and Figure S5A-D**). Across the immune compartment, we observed reproducible increases in follicular helper T cells (CD4_Tfh), macrophages (Macro_TREM2 and Macro_SPP1), cytotoxic CD8+ T cells (CD8_Tex_GZMK), and cycling germinal center B cells (GCB-cycling), accompanied by reductions in interferon-responsive T cells (CD4_T-ISG and CD8_T-ISG). Within the stromal and vascular compartments, progenitor and inflammatory cancer-associated fibroblasts (CAF-prog and iCAF_MMP1) were enriched in treated tumors, whereas myofibroblasts and tip-like endothelial cells (Endo-tip) were consistently depleted.

The temporal dimension of remodeling revealed additional complexity. TIME dynamics differed markedly after a single ICI cycle compared with two or more cycles (**Figure S6A-B**). Early (one-cycle) remodeling was characterized by an increase in CD4_Tfh cells and a decrease in γδ T cells, alongside transient monocyte depletion (**Figures S6A-B**). At this stage, SKCM tumors exhibited a reduction in immunosuppressive macrophages (Macro_LYE1 and Macro_C1QC), whereas ER+ BRCA and TNBC displayed the opposite trend. Later (two-plus-cycle) remodeling involved myeloid enrichment, most evident in HNSCC receiving anti-PD-1 monotherapy and NSCLC treated with anti-PD-1 plus chemotherapy, while combination therapy in HNSCC led to depletion of cDC2 and mregDC populations. In the B-cell compartment, a transient increase in naïve B cells was detected after the first cycle but was not maintained in TNBC and HNSCC after extended treatment (**Figure S6C**). Parallel stromal remodeling featured early expansion of endothelial (Endo-lymphatic and Endo-capillary) and iCAF_MMP1⁺ fibroblasts, coupled with progressive loss of smooth-muscle cells (SMC), myofibroblasts, and pericytes over time (**Figure S6D**).

We next examined whether cell-state dynamics correlated with treatment efficacy (**Figures S7-S8**). Responders showed a sustained increase in exhausted CD8⁺ T cells after two or more cycles, accompanied by depletion of effector-like CD8 subsets (**Figure S7A**). In contrast, immunosuppressive macrophages (Macro_LYE1, TREM2, C1QC) decreased in SKCM responders after one cycle but persisted or expanded in non-responders, particularly in ER+ BRCA (**Figure S7B**). B-cell remodeling also differed by outcome: non-responders displayed an attenuated, often non-significant, decrease in activated and memory subsets (**Figure S8A**). Among non-immune lineages, responders consistently exhibited reduction of tip-like endothelial cells (Endo-tip), suggesting vascular normalization as a potential feature of effective ICI response (**Figure S8B**).

These results outline the TIME changes during ICI therapy. The process involves coordinated activation of helper and cytotoxic T cells, expansion of selected macrophage populations and B cell subsets, and measurable adjustments in fibroblast and endothelial compartments. Together, these shifts form a pattern of immune remodeling that appears linked to treatment outcome.

### Four distinct TIME subtypes capture coordinated cellular organisation

Individual immune-subtype changes typically represented only small fractions of total immune infiltration, making global interpretation difficult. To identify coordinated cellular programs that shape the TIME, we applied non-negative matrix factorization (NMF) to the immune-cell composition profiles from 418 samples, reserving our own melanoma cohort for independent validation (**Figure 3A**). NMF rank survey metrics identified four as the optimal number of programs (**Figure S9A)**, which captured major axes of microevironmental heterogeneity. Each tumor was assigned to a subtype based on its NMF membership scores (**Table S4**), with clear separation demonstrated by the distinct loading patterns of immune-cell subtypes across the programs (**Figure 3B**). Because all tumors from the HCC_Guo cohort were mapped to a single subtype, they were excluded from downstream analyses to avoid study-specific bias. NMF classifications showed no significant difference between CD45+ sorted and entire TME samples in HNSCC (with a sufficient number of samples), indicating robustness to sample processing protocol (P=0.395, Fisher’s exact test; **Table S4**).

**Figure 3.**
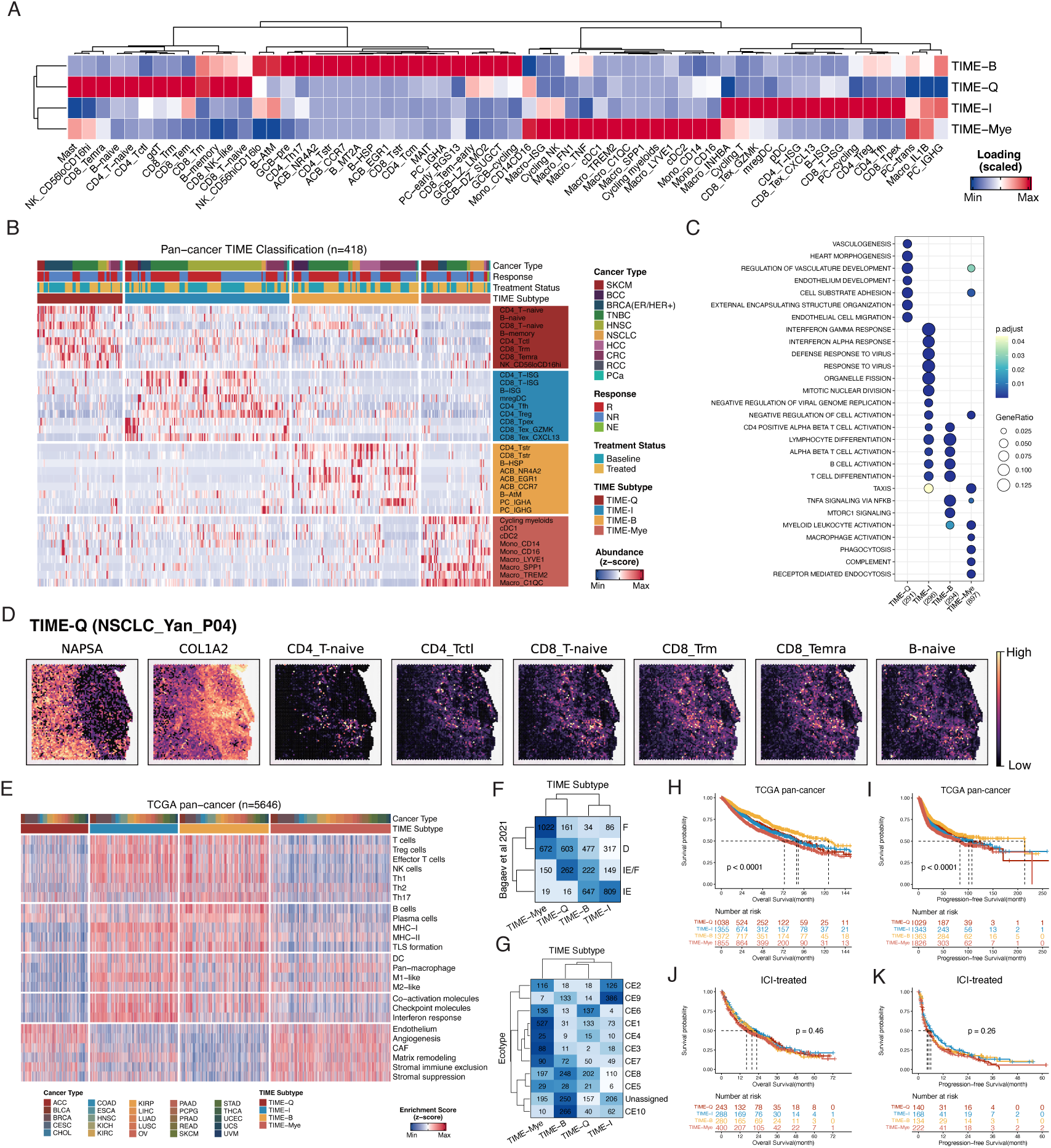
Four distinct TIME subtypes define coordinated immune–stromal organization. **(A)** NMF-based cellular programs derived from immune cell abundance profiles. Columns represent immune subtypes and rows indicate loading values for each program. (B) NMF clustering identifies four TIME subtypes, each enriched for distinct immune states. Heatmap colors indicate the relative contribution of each cell type; representative cellular modules for each subtype are shown on the right. **(C)** Dot plot showing functional pathways enriched for markers of the four TIME subtypes. **(D)** Spatial feature plots displaying gene expression and deconvoluted cell-type abundance in a representative TIME-Q tumor from the NSCLC_Yan cohort. **(E)** Heatmap of curated TME signature enrichment scores across TIME subtypes. **(F)** Comparison of TIME subtypes with conserved pan-cancer TME classes (Bagaev et al.). **(G)** Comparison of TIME subtypes with carcinoma ecotypes (Luca et al.). **(H–I)** Kaplan–Meier curves showing overall survival (OS) and progression-free survival (PFS) in TCGA pan-cancer bulk RNA-seq data. **(J–K)** Kaplan–Meier curves showing OS and PFS in aggregated bulk transcriptomic data from ICI-treated cohorts.

TIME-I (immune-inflamed) tumors exhibit high infiltration of both adaptive immune cells, including interferon-responsive and CXCL13^+^ T cells, B cells, and plasma cells, alongside inflammatory macrophages (**Figure 3A-B**). Functional enrichment confirmed activation of immune-effector and interferon pathways (**Figure 3C**), although concurrent accumulation of exhausted T cells and regulatory DCs suggested a mixed state characterized by both immune engagement and regulatory suppression mechanisms. TIME-B (B-cell-enriched) tumors are characterized by abundant activated and atypical memory B cells and plasma cells, accompanied by CD4+ central memory T cells and type 17 T cells (Th17), reflecting active humoral immunity and potential tertiary lymphoid structure (TLS) formation. TIME-Mye (myeloid-enriched) tumors are dominated by various myeloid cell populations, including macrophages, monocytes, and DCs, indicative of an immunosuppressive, myeloid-rich environment. TIME-Q (immune quiescent) tumors are defined by early-stage T and B cells and non-exhausted cytotoxic T cells. Spatial transcripromics analysis revealed that these populations are primarily localized to stromal areas along tumor boundaries, resembling the immune-excluded phenotype previously described in liver cancer (**Figure 3D**).^43^

Beyond immune composition, the TIME subtypes reflected broader microenvironmental organization. TIME-Q and TIME-Mye displayed higher stromovascular content compared to TIME-B and TIME-I (**Figure S9B**). Inflammatory and antigen-presenting CAF populations were selectively enriched in TIME-B tumors (**Figure S9C**), supporting the concept of coordinated pro-inflammation immune-stromal interactions within this subtype. Curated TME signature analysis further validated these functional distinctions: TIME-I showed the highest scores for cytotoxicity, co-activation molecules, checkpoint molecules, and interferon response, whereas TIME-B demonstrated elevated Th17 and B/plasma cell signatures (**Figure S9D-E; Table S5)**. Cell-cell communication analysis suggested that the interactions among T cells and NK cells in TIME-Q were substantially lower compared to TIME-I and TIME-B (**Figure S9F**). To assess the robustness of our classification, we scaled the relative proportions of our melanoma validation cohort and recovered the NMF weight for sample assignment. The distribution pattern persisted compared to the atlas, suggesting that the classification was indeed based on inherent biological variation between individuals (**Figure S9G; Table S4**).

To relate our TIME subtypes to previously established TME frameworks, we projected them onto TCGA pan-cancer bulk RNA-seq data (**Methods**; **Figure 3E; Table S6)** and compared our subtypes to the conserved subtypes^9^ (**Figure 3F**), pan-cancer immune subtypes^10^ (**Figure S10A**), and carcinoma ecotypes (CE)^38^ (**Figure 3G**). The comparison revealed both shared features and key distinctions among systems. TIME-I overlapped most strongly with the ‘immune-enriched’, ‘IFN-gamma dominant subtype’ and CE9, reflecting its high immune lymphocyte infiltration and interferon signaling. TIME-B showed partial similarity to the ‘immune-enriched’ and proinflammatory CE10 clusters, consistent with its activated B-cell and plasma-cell composition. TIME-Q and TIME-Mye aligned more closely with stromal-enriched and lymphocyte-depleted subtypes such as ‘fibrotic/depleted’ and CE6. TIME-Mye also mapped to ‘wound healing’ and ‘lymphocyte-deficient’, and the survival-inferior CE1-CE4 ecotypes, indicating that it captures features from both lymphocyte-desert and myeloid-suppressive microenvironments associated with poor progrnosis. Consistent with these biological patterns, TIME-I and TIME-B tumors exhibited the highest tumor mutational burden (TMB), T-cell receptor (TCR) richness, and tumor purity (**Figure S10B**), supporting their classification as immunologically active subtypes.^44^ Projecting the single-cell-derived classification onto TCGA bulk transcriptomes also confirmed prognostic relevance: across cancer types, TIME-B showed significantly longer overall and progression-free survival, whereas TIME-Mye was consistently associated with the worst outcomes (**Figure 3H-I**).

To test whether baseline immunotypes also predict outcomes under checkpoint blockade, we transferred the classification to a pan-cancer compendium of 1,988 pre-treatment RNA-seq samples from ICI-treated patients (**Table S7**).^45^ Each sample was assigned a TIME label by deconvolution (**Figure S10C**). Baseline response rates were higher for TIME-I (36.9%) and TIME-B (31.1%) than for TIME-Q (22.3%) and TIME-Mye (26.0%) (P = 0.04; **Figure S10D**). In contrast to TCGA, however, these pre-treatment immunotypes did not significantly separate overall or progression-free survival in the ICI cohorts (**Figure 3J-K**). This divergence indicates that while the subtypes capture intrinsic tumor immunobiology and long-term prognosis, a static baseline label is insufficient once therapy perturbs the microenvironment. These results motivate the shift to longitudinal TIME dynamics as the relevant signal for immunotherapy sensitivity and resistance.

### TIME transitions were associated with treatment efficacy

Longitudinal pairing of biopsies across our pan-cancer atlas enabled a systematic analysis of TIME remodeling during ICI therapy. Among 187 tumors, 57.8% (N=108) maintained their baseline subtype through treatment, significantly more than expected by random chance (P < 0.0001, exact binomial test; **Figure 4A**), while the remaining 42.2% (N=79) underwent subtype transitions. The consistency of these transition rates across treatment dosage suggests these shifts represent sustained, treatment-associated reprogramming rather than transient sampling variability (P=0.437, Fisher’s exact test; **Figure 4B; Table S7**).

**Figure 4.**
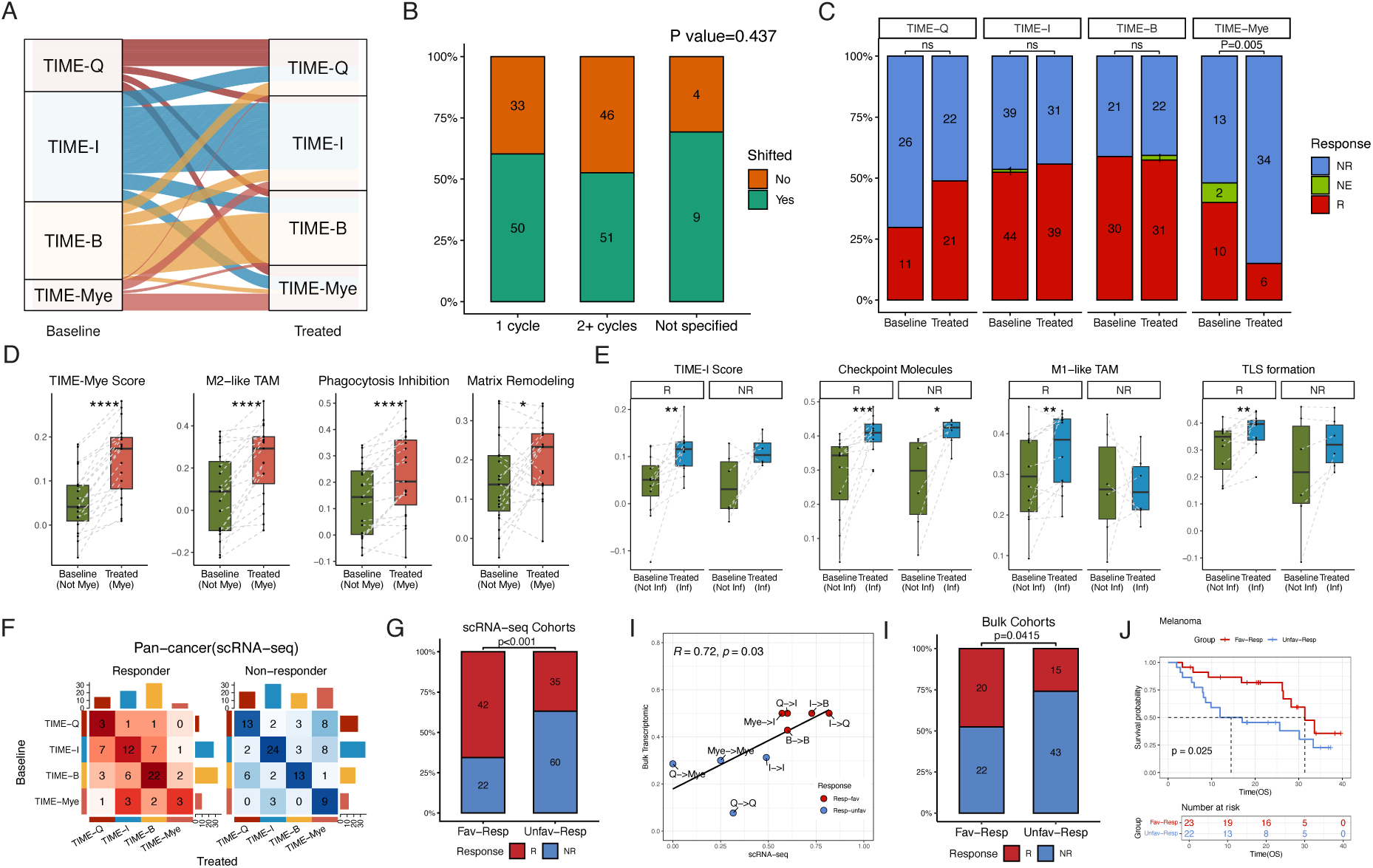
TIME transitions during ICI therapy reflect and predict treatment efficacy. **(A)** Alluvial diagram showing TIME subtype transitions between baseline and post-treatment biopsies in ICI-treated tumors. **(B)** Bar plot summarizing the proportion of tumors undergoing subtype transitions across different treatment durations. **(C)** Distribution of responders and non-responders among baseline and post-treatment TIME subtypes. **(D)** Boxplots comparing immune-suppression–related signatures (M2-like macrophages, phagocytosis inhibition, matrix remodeling) between baseline and post-treatment samples in tumors that transitioned toward the TIME-Mye subtype. Statistical testing by paired Wilcoxon signed-rank test. **(E)** Boxplots showing activation of anti-tumor programs (TIME-I signature, checkpoint-molecule expression, M1-like macrophage polarization, TLS formation) in tumors transitioning toward the TIME-I subtype, stratified by standardized response. Paired Wilcoxon signed-rank test as in (D). **(F)** Heatmap displaying the frequency of subtype transitions in responders and non-responders. **(G)** Bar plot comparing response rates between response-favorable and response-unfavorable transition groups defined in the single-cell cohort. Statistical testing by Chi-squared test. **(H)** Scatter plot showing correlation of transition-specific response probabilities between single-cell and bulk RNA-seq paired datasets (Spearman correlation). **(I)** Bar plot comparing response rates between response-favorable and response-unfavorable transitions in paired bulk RNA-seq data. Statistical testing by Chi-squared test. **(J)** Kaplan–Meier curves showing overall survival for response-favorable and response-unfavorable transition groups in the melanoma bulk cohort (one-sided log-rank test). *p < 0.05, **p < 0.01, ***p < 0.001, ****p < 0.0001

Treatment outcome was closely tied to these subtype trajectories. Non-responders were enriched for treated TIME-Mye tumors (P = 0.002, Chi-squared test; **Figure 4C**), which upregulated immunosuppressive programs, such as M2-like macrophages, phagocytosis inhibition, and matrix remodeling (**Figure 4D**). This pattern suggests acquisition of a myeloid-dominant, therapy-resistant phenotype. In contrast, responders whose tumors transitioned towards the TIME-I subtype demonstrated a coordinated upregulation of anti-tumor pathways, including enhanced TIME-I signature, checkpoint-molecules expression, M1-like macrophage activity, and TLS formation (**Figure 4E**).

Transition direction also differed by outcome. For instance, among patients with baseline TIME-I tumors, responders were significantly more likely to transition toward the TIME-B or TIME-Q subtypes, whereas non-responders showed higher rates of transitions toward the response-unfavorable TIME-Mye subtype (P = 0.007, Fisher’s exact test; **Figure 4F**). Within TIME-Q tumors, responders exhibited significantly higher B-cell infiltration compared to non-responders **(Figure S11A),** suggesting crucial B-cell-centric immune activation for positive therapeutic outcomes. Favorable trajectories varied by cancer type: responders with SKCM and HNSCC preferentially switched to TIME-Q, whereas responsive TNBC tumors were more likely to transition to TIME-B **(Figure S11B).**

Given that subtype transitions were directly correlated with treatment outcome, we posit that TIME dynamics provide a more informative representation of treatment efficacy. We classified the observed transitions based on clinical outcomes into response-favorable and response-unfavorable categories (**Figure 4F**). The response-favorable category includes the transitions: TIME-B/I→TIME-B and TIME-Q/Mye→TIME-I. The response-unfavorable category includes the transitions: maintenance of TIME-Q, TIME-Mye, or TIME-I states, TIME-Q→TIME-Mye, and TIME-B→TIME-Q. By definition, patients in the response favorable group exhibited higher objective response rates than those in the response-unfavorable group (P < 0.001, Chi-squared test, **Figure 4G**).

To evaluate the reproducibility of these transition patterns in independent datasets, we next applied the same response-favorable and response-unfavorable classification to bulk RNA-seq cohorts with paired biopsies from 94 ICI-treated patients across multiple cancer types. Using the transition definitions derived from the single-cell atlas, we computed response probabilities for each transition pattern in the bulk data. These probabilities showed strong concordance between the single-cell and bulk cohorts (R = 0.72, P = 0.03; **Figure 4H**), confirming that the response-associated dynamics captured by our classification are robust across data modalities. Similarly, patients undergoing response-favorable transitions displayed a higher likelihood of clinical response than those with response-unfavorable transitions (P = 0.0415, Chi-squared test; **Figure 4I**). Furthermore, melanoma patients with response-favorable transitions had longer overall survival than those with response-unfavorable dynamics (HR: 2.24, 95% CI 0.98- 5.15, P = 0.056; **Figure 4J**).

Together, these results demonstrated that TIME transitions detected from paired samples reliably reflect treatment efficacy and clinical outcome. Monitoring such dynamic remodeling, even after the first treatment cycle, thus provides a sensitive and generalizable surrogate for therapeutic response.

### Baseline Features Shape TIME Trajectories and Predict ICI Response

Given the close association between TIME transitions and treatment efficacy, we hypothesized that the TIME trajectory under ICI therapy is influenced by its baseline characteristics. In this view, pre-existing features could determine transition fate and, in turn, treatment outcome. To test this hypothesis, we performed differential expression analysis on pseudobulk profiles of baseline TIME-I tumors that underwent distinct trajectories. Focusing on the TIME-I→TIME-B/Q (response-favorable) versus TIME-I→TIME-Mye (response-unfavorable) transitions, we identified distinct baseline programs associated with transition fate (**Figure 5A; Table S8**). From these gene sets, we derived a composite transition score quantifying the likelihood of a favorable trajectory. The score was significantly higher in TIME-I→TIME-B/Q tumors than in TIME-I→TIME-Mye tumors (**Figure 5B**), and was also elevated in clinical responders versus non-responders **(Figure 5C)**, representing a predictive metric of both TIME remodeling and clinical outcome.

**Figure 5.**
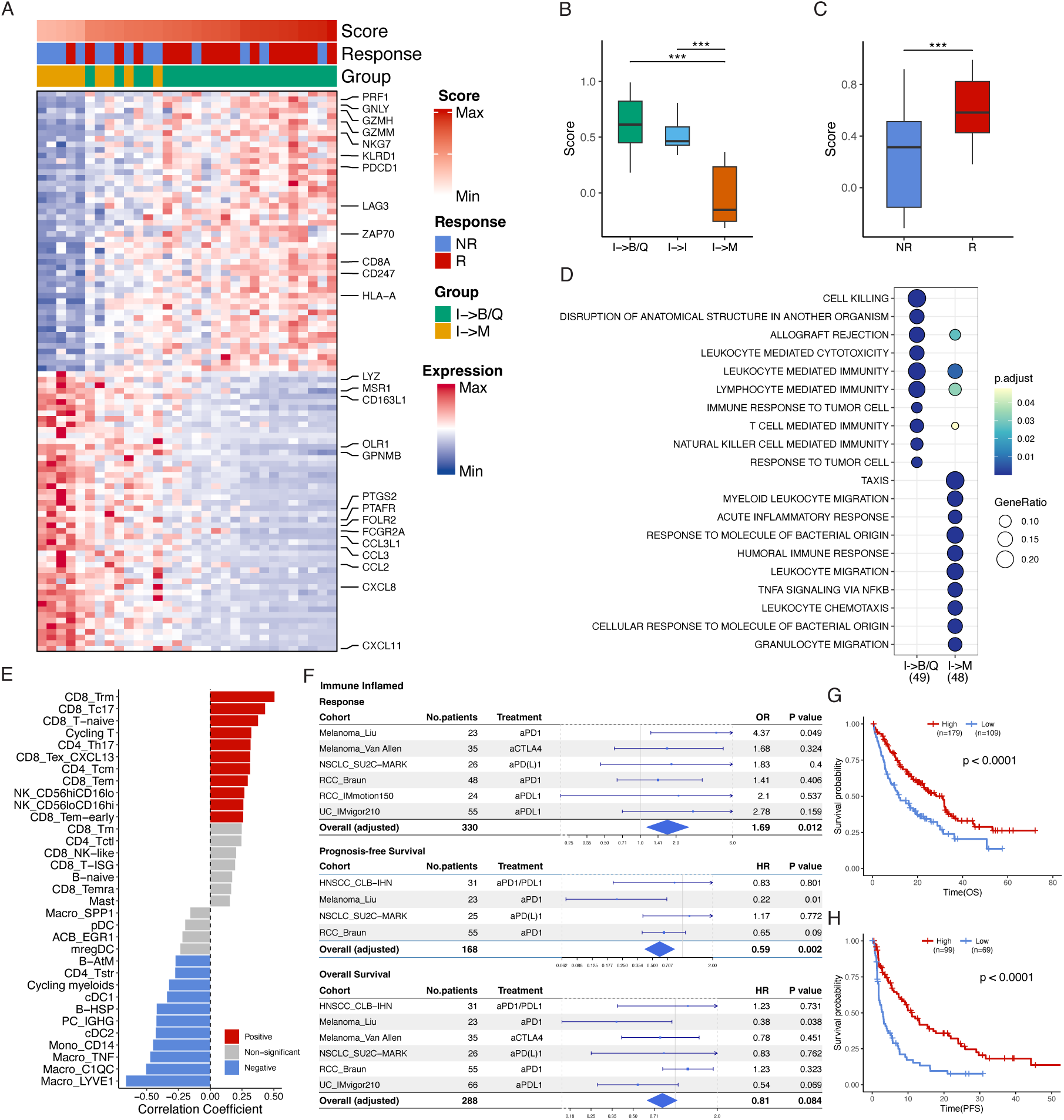
Baseline features of inflamed tumors predict immune evolution and clinical outcome. **(A**) Heatmap of the top 50 upregulated and 50 downregulated genes distinguishing baseline TIME-I tumors that transitioned toward TIME-B/Q versus TIME-Mye subtypes during treatment. **(B)** Boxplots comparing transition scores across TIME-I tumors with distinct transition fates. Two-tailed unpaired Wilcoxon tests. **(C)** Boxplots comparing transition scores between responders and non-responders among baseline TIME-I tumors. Two-tailed unpaired Wilcoxon tests. **(D)** Dot plot showing enriched pathways for genes defining response-favorable (TIME-I → TIME-B/Q) and response-unfavorable (TIME-I → TIME-Mye) transitions. **(E)** Bar plot showing Spearman correlation coefficients between transition scores and immune-cell subtype abundance in baseline TIME-I tumors. **(F)** Forest plots summarizing predictive performance of the transition score for treatment response, progression-free survival (PFS), and overall survival (OS) across baseline bulk transcriptomic ICI cohorts. **(G–H)** Kaplan–Meier curves for PFS and OS comparing patients with high versus low baseline transition scores. Two-sided log-rank tests.

Baseline profiles of response-favorable tumors were enriched for cytotoxic and effector T-cell genes such as PRF1, CD8A, and GZMH, together with immune-checkpoint molecules PDCD1 and LAG3, indicating a primed yet regulated immune state. Pathway analysis confirmed enrichment for leukocyte-mediated cytotoxicity, T-cell activation, and NK cell responses (**Figure 5D**). In contrast, tumors destined for response-unfavorable transitions displayed dominant myeloid programs, including M2-like macrophage markers MRC1 and CD163L1 and pro-inflammatory cytokines IL1B and CXCL8, consistent with chronic rather than productive inflammation. These genes were enriched for myeloid migration, chemotaxis, and matrix remodeling pathways. Correlation of the transition score with cell-type composition revealed strong positive associations with cytotoxic CD8⁺ T cells, Th17 cells, and NK cells, and negative associations with suppressive macrophages, CD14⁺ monocytes, cDC2, and IGHG⁺ plasma cells (**Figure 5E**). Among stromal populations, iCAF, CAF-prog, and CAF_SFRP2 were enriched in TIME-I→TIME-B/Q tumors, whereas antigen-presenting stromal cells predominated in TIME-I→TIME-Mye trajectories (**Figure S12A**).

Importantly, validation in independent baseline bulk transcriptomic cohorts of ICI-treated patients confirmed that the transition score, computed from pretreatment samples alone, robustly predicts clinical outcome (**Figure 5F**). Across 1,988 patients from multiple cancer types, 484 tumors were classified as TIME-I at baseline and thus eligible for score-based stratification. Higher transition scores were significantly associated with greater response probability (OR = 1.69, 95% CI: 1.13-2.56, P = 0.012), longer progression-free survival (HR = 0.59, 95% CI 0.42-0.83, P = 0.002), and a trend toward improved overall survival (HR = 0.81, 95% CI 0.63-1.03, P = 0.084). Kaplan-Meier analysis further showed that patients with high baseline transition scores had markedly longer PFS and OS (both P < 0.001; **Figure 5G-H**).

Parallel analysis of TIME-Mye baseline tumors revealed additional determinants of therapeutic success. Responders were more likely to transition from TIME-Mye to TIME-I, while non-responders tended to remain myeloid-stable (P = 0.07, Fisher’s exact test). Tumors predisposed to favorable TIME-Mye→TIME-I transitions expressed CXCL9 and atypical B-cell marker FCRL4, reflecting preserved capacity for lymphocyte recruitment and microenvironmental repolarization. In contrast, non-responders exhibited sustained myeloid programs marked by CD33, PKIB, and LRP3, consistent with stable immunosuppressive polarization (**Figure S12B; Table S8**). Validation in bulk transcriptomic cohorts reproduced the predictive performance of the transition score across multiple clinical endpoints: for treatment response (OR = 3.05, 95% CI: 1.56-6.19, P = 0.001), PFS (OR = 0.41, 95% CI: 0.21-0.81, P = 0.01), and OS prediction (hazard ratio = 0.68, 95% CI: 0.46-1.03, P = 0.066) (**Figure S12C**).

Comparing TIME-I- and TIME-Mye-based analyses revealed both conserved and context-dependent determinants of remodeling. Conserved features included a positive correlation with CD8_Tex_CXCL13 and a negative correlation with Macro_C1QC across both transition types (**Figure S12D**). Context-dependent immune features were also identified: B-AtM, stress-associated CD4-str, B-HSP, and PC_IGHG correlated positively with the TIME-Mye transition score and negatively with the TIME-I transition score. Similarly, conserved stromal features included enrichment of iCAF_MMP1 in response-favorable transitions and CAF-ap/capillary-like endothelial cells in response-unfavorable transitions across both TIME subtypes (**Figure S12E)**. In a context-dependent manner, CAF_prog, iCAF_IL6, and CAF_SFRP2 were enriched in response-favorable transitions of TIME-I but in response-unfavorable transitions of TIME-Mye.

In summary, a baseline-derived transition score quantifies the intrinsic potential of a tumor’s microenvironment to evolve toward an immune-active state under therapy. These findings reveal conserved and context-specific cellular programs that pre-determine the direction and efficacy of TIME remodeling during ICI treatment.

## Discussion

Studies that followed patients with sequential biopsies have been invaluable for revealing the contours of this remodeling in cancers such as melanoma, lung, head and neck, breast, liver, and colorectal cancers.^15,20–23,25–28,30,46,47^ Yet these prior analyses have been constrained by sample size. Cohorts of 10 to 20 paired cases are large for a single-cell study, but they are not powered to resolve modest, clinically relevant effects. Single-cell studies aimed at clinical translation require the statistical advantages of larger, harmonized compendia. Our design follows that blueprint by integrating longitudinal single-cell datasets, imposing a unified two-level reference-guided annotation, and testing findings across bulk RNA cohorts. Taken together, this framework allows us to show that remodeling of the TIME carries more prognostic value than static baseline immunotypes.

Our reference-guided deep-phenotyping framework to align cell types with comprehensive pan-cancer reference data across studies achieved high-resolution annotations while circumventing the over-reliance on integration pipelines that lack consensus benchmarks and reproducibility.^5,48,49^ Our meta-analysis identified consistent compositional changes across studies, including increased frequencies of follicular helper CD4 T cells, TREM2-positive macrophages, and inflammatory cancer-associated fibroblasts, together with reduced proportions of interferon-responsive T cells and tip-like endothelial cells during treatment. Subgroup analyses showed that these shifts were not uniform across patients but varied with treatment dose, regimen, and clinical response, highlighting their relevance to therapy-associated remodeling.

Based on compositional data, we identified four reproducible TIME subtypes that capture coordinated cellular programs. Projection to TCGA data showed that these subtypes align with but are not redundant to established classifications.^9,10,38^ The distinct TIME-B from immune-enriched subtypes exhibited superior prognostic stratification on TCGA primary tumors. In contrast, in a large compendium of pre-treatment biopsies from ICI-treated patients, we found that this baseline classification alone was not associated with clinical outcome. Previously, Bagaev et al. reported TIME subtype dynamics on bulk RNA-seq data in melanoma.^9^ In our pan-cancer scRNA-seq atlas, we confirmed that a substantial proportion of tumors underwent TIME subtype transitions during the treatment. While the shift rate was constant over time, the specific transitions showed an association with clinical response. Specifically, a shift toward a myeloid-dominant immunosuppressive state correlated with poorer outcomes, while a shift toward a B-cell-centered, immune-activating state was associated with improved response across single-cell and bulk datasets. These patterns place the B cell and myeloid modules within a coherent dynamic framework and reinforce their central influence on how the TIME evolves under therapy.^50,51^ We demonstrate that TIME subtype transition represents a conserved property of immune reinvigoration. Therefore, it could serve as a valuable surrogate biomarker that complements other emerging monitoring strategies, such as circulating T cells and tumor DNA.^52,53^

We derived a transition score from baseline inflamed tumors that predicted polarization toward divergent TIME fates: a B cell-centric state versus a myeloid-suppressed state. This score successfully predicted treatment response and survival in independent bulk RNA-seq cohorts, capturing cellular features that may prime the tumor for its subsequent immune trajectory. Although the constituent genes reflect well-established immune features, such as effector T cells and immunosuppressive myeloid populations, our results provide novel insights by framing them as a cohesive predictor of immune trajectory. For pre-treatment stratification, this score adds a dynamic dimension to conventional baseline biomarkers such as PD-L1 expression and tumor mutation burden, thereby potentially facilitating more informed clinical decision-making.^54,55^

Several limitations merit attention. Despite harmonization, residual batch and sampling biases are possible, including differences in biopsy timing, lesion site, prior lines, and concurrent therapies. Response categories and survival endpoints are not uniform across cohorts. Spatial information is limited outside of the cohorts that include Visium or multiplex imaging. Although validation across single-cell and bulk modalities increases confidence, prospective studies that embed pre-specified dynamic rules and decision thresholds are necessary to test clinical utility. Finally, while the transition score performed consistently across multiple cancers, cutoffs for clinical decision-making will require calibration by indication.

In summary, by scaling sequentially paired single-cell profiling through rigorous integration and cross-modal validation, we showed that dynamic TIME transitions outperform static baseline labels for monitoring response, and that a baseline-derived transition score predicts benefit in independent bulk cohorts. The field has reached a point where cellular diversity is well cataloged; the path to clinical impact requires larger, harmonized, longitudinal resources and analysis plans built for power and reproducibility. Our study provides one such template and supports a broader shift toward meta-analytic single-cell oncology that can sustain reliable, treatment-guiding inferences at the level patients and clinicians need.

## Methods

### Single-cell RNA-seq data generation

Patients with advanced melanoma who were to receive immunotherapy with anti-PD-1 antibodies alone or in combination with anti-CTLA4 antibodies from Departments of Translational Skin Cancer Research and Dermatology were included in the study upon written informed consent (ethics vote numbers 16-6784-BO und 16-7097-BO). Tumor tissue biopsies from melanoma lesions were obtained on the day of therapy initiation (day 0) and 7 days thereafter (day 7). Fresh biopsies were immediately processed after excision. The generation of the single cell suspension was performed according to the tumor dissociation protocol from Miltenyi Biotec (Bergisch Gladbach, Germany). Briefly, the tissue biopsy was cut into small cubes of 2 mm and then minced with the gentleMACS program “h_tumor_01” and two times “h_tumor_02” with 30 min incubation time after each mince. During the incubation, the sample was digested with an enzyme cocktail mix consisting of 200 µl Enzyme H, 100 µl Enzyme R and 25 µl Enzyme A (catalogue #130-095-929 Miltenyi Biotec). The cell suspension was reconstituted in PBS with 0.05% BSA and washed two to three times, before passing through a 100 µm cell strainer. Subsequently, the cells were processed with the Single Cell 5’ Library & Gel Bead Kit v1.1 (catalogue #1000165, 10xGenomics, Pleasanton, CA, USA) according to the manufacturer’s protocol. As input for the 10x protocol, 10,000 cells from the single cell suspension, at a concentration of 1000 cells/µl. The resulting gene expression library was sequenced at genomics core facility of German Cancer Center (DKFZ), Heidelberg, Germany. The sequencing data were aligned with the GRCh38 human reference genome and quantified using Cell Ranger (version 3.0, 10x Genomics Inc).

### Single-cell RNA-seq data curation and standardization

Through an extensive literature review up to Mar 2025, we identified public scRNA-seq datasets across multiple cancer types with a specific focus on samples collected at the baseline and after the initiation of the ICI therapy (**Table S1**).

Acknowledging the diverse response metrics used across neoadjuvant scRNA-seq studies, including RECIST, pathological assessment, PSA tests, T cell expansion, and early radiological evaluations, we harmonized responder classification for comparative analysis. Patients were designated as responders if they met any of the following criteria: achieving a complete or major pathological response, demonstrating a complete or partial response by RECIST, or being a pathological non-responder who nonetheless showed a RECIST response. In cohorts where response was primarily assessed via T cell expansion or blood prostate-specific antigen (PSA) levels, the original study’s classification was adopted. Notably, consistent with the threshold used in BRCA_Bassez cohort from the same group, for the HNSCC_Franken cohort, patients with > 30 T cell expanded clonotypes were classified as responders.

Given the wide sampling interval spanning 7 to 121 days from the first dose of immunotherapy, all samples were classified as either baseline (pre-treatment) or treated (on/post-treatment) status.

### Single-cell RNA-seq data preprocessing

Newly generated single-cell sequencing data were aligned with the GRCh38 human reference genome and quantified using Cell Ranger (version 3.0, 10x Genomics Inc). The quality of cells was assessed based on two metrics: (1) The number of detected genes per cell; and (2) The proportion of mitochondrial gene counts per cell. Specifically, cells with detected genes fewer than 400 or greater than 8000, and mitochondrial unique molecular identifier (UMI) count percentage larger than 20% were filtered out. The genes detected in more than 5 cells were retained. To remove the potential doublets, scDblFinder^11^ and DoubletFinder^12^ were used for each sequencing library with the expected doublet rate set to be 0.08, and cells identified as doublets by both methods were further filtered out. For each dataset, gene symbols were converted into a common dictionary with STACAS:::standardizeGeneSymbols()^13^. After the initial quality control and cell annotation, sample-level quality control was applied to remove samples with fewer than 200 non-malignant cells.

### Hierarchical cell annotation

The hierarchical annotation framework facilitates consensus and flexible cell type assignment at two levels, i.e., main cell types and fine-granular subtypes.

First level annotation was performed within individual datasets. SingleR^14^ with the BlueprintEncodeData^15,16^ reference data was applied to predict cell types. The annotation was refined by scGate^56^ with a pre-defined multi-classifier (TME_HiRes) and Seurat V5 Louvain clustering^57^ (default resolution) on corrected embeddings by Harmony^58^. Cells labeled as ‘multi-label’ and clusters with gene expression from multiple cell lineages were identified as doublets. Due to the technical limitations of 10X sequencing, the neutrophils were also excluded from subsequent analysis. The raw annotation from this step was improved by the following level 2 annotation and the final cross-dataset dataset integration. Malignant cells were identified based on inferred copy number aberration (CNA) by infercnv package (see **Malignant cell identification** section).

Second level annotations were designed to cater to various cellular lineages, and consisted of two primary steps, i.e., reference preparation and hierarchical annotation.

For reference preparation, reference data for T-cells, NK cells, myeloid cells, and CAFs were retrieved from the pan-cancer atlas studies^3,4,6,8,20,21^, loaded in R as *SingleCellExperiment* objects, and processed individually. The objects were split by cancer type or cohort as lists, normalized with *logNormCounts*(), and further aggregated into pseudo-bulk samples *aggregateReference*() by the fine-granular level labels from original studies. Additionally, to capture the diversity of main components, data for T cells, myeloid cells, and B lymphocytes were also aggregated at the broad level (e.g., CD4+ T cells, CD8+ T cells, cDC, macrophages, B cells, plasma cells) to ensure accurate reclassification. Due to the relatively lower abundance, the reference data of NK cells was aggregated only at the main level (CD56^high^CD16^low^, CD56^high^CD16^high^, and CD56^low^CD16^high^).

For hierarchical annotation, immune cells were initially re-classified into main cell types within each lineage by SingleR based on the pan-cancer reference at the main level. For the T and NK cell group, this included CD4+ T cells, CD8+ T cells, and NK cells. For the myeloid cells group, this included cDCs, monocytes, and macrophages. For the B lymphocytes, this included B cells, GC B cells, and plasma cells. Due to the distinct marker expression, the scGate was applied to separate cells from the T and NK cells group, with *KLRD1* and *NCAM1* served as positive markers, and CD3D as a negative marker for NK cells, and *CD3D*, *TRGC2*, *TRDV2*, *TRGV9*, *TRGV10*, and *TRDC* as positive markers, and CD8A and CD4 as negative markers for the gamma delta T cells (gdT). After separating the NK and gdT, the remaining CD4+/CD8+ T cells underwent further ‘gating’ by the generic CD4T and CD8T models from scGate. Cells with overlapped labels were identified by SingleR, based on an aggregated pan-cancer reference at the main level. Similar procedures of reassignment at the first level were conducted on myeloid cells and B cells. The immune cells were then classified within each main cell type by SingleR with pan-cancer multiple reference at the fine-granular level. Non-immune cells including endothelial cells and fibroblasts cells were annotated directly to the fine-granular level by using SingleR with corresponding reference. After prediction, the labels were adjusted according to the extensively updated markers.

Finally, as a manual adjustment and quality control step for the final annotation, we performed cross-dataset integration within each cell lineage by Seurat integration. This step not only adjusted the annotation at different levels but was also instrumental for excluding problematic clusters. Clusters with high expression of proliferation-associated such as MKI67 and TOP2A were identified as cycling cell states. Clusters with clear expression of gene markers from multiple cell lineages and with high expression of ribosomal genes were identified as doublets and low-quality cells then were removed.

### Malignant cell identification

The malignant cells detection was performed by using the inferCNV^59^ package patient by patient. This process infers large-scale chromosomal copy number variations (CNVs) from single-cell RNA-sequencing data. First, the data were subsetted by patient, including epithelial cells (which are considered the origin of the tumor), and a reference population of various non-malignant cell types like fibroblasts and monocytes. The CNV states were predicted with a three-state CNV model (HMM_type=’i3’). Cells with both average proportion of CNV-affected chromosomes (proportion_cnv_avg) and the average number of CNV events (has_cnv_avg) above the 90th percentile were classified as malignant.

### Cell type composition

Samples with no less than 100 non-epithelial/malignant cells and no less than 50 immune cells were included in compositional analysis. Cell counts from of each subtype were divided by the total immune/non-immune cells.

### Meta-analysis of inter-cohort compositional analysis

To assess the consistency of cell type abundance changes between baseline and treated states and account for across-study variability, we conducted a two-stage meta-analysis for each cell type: First, within each individual cohort, we used the Wilcoxon rank-sum test (Mann-Whitney U test) to compare the abundance of each cell type between baseline and treated status. For each comparison, the effect size was calculated as the rank-biserial correlation 𝑟, a non-parametric measure of association, using the rcompanion R package. We estimated the 95% confidence interval for 𝑟 via 1000 bootstrap replicates. The standard error 𝑆𝐸 for each effect size was then approximated from its 95% confidence interval. Cohort-cell type combinations were excluded if they had fewer than two groups for comparison or insufficient number of samples. Second, to summarize results for each cell type across all cohorts, we performed a random-effects meta-analysis using the metafor R package. This model pooled the per-cohort 𝑟 as the effect size estimates and used their corresponding 𝑆𝐸 for inverse-variance weighting. We employed the Restricted Maximum Likelihood (REML) estimator to compute the pooled summary effect, its 95% confidence interval, and the p-value.

### Gene set enrichment analysis

Gene set enrichment for scoring at bulk (including pseudobulk) and single cell level was conducted by using ssGSEA method from GSVA package^60^ and *AddModuleScore()* from Seurat^57^, individually.

### Differential expression analysis and over-representation enrichment analysis

The differential expression (DE) analysis was performed by using *FindAllMarkers* () function from the Seruat package based on the Wilcoxon rank-sum test. Mitochondrial genes and ribosomal genes were removed before DE analysis. In pseudobulk DE analysis, minimum fraction of detection of 0.9 was add as filtering criteria. The functional enrichment analysis was performed by using the clusterProfiler package.

### Non-negative matrix factorization (NMF) analysis for identifying tissue-specific cellular programs

With the compositional data as matrix 𝑉 (cell type × sample), we performed NMF by using R package NMF across samples from both pre- and -on treatment time points to identify tissue-specific programs within the TME. With n samples and m cell types, V is a non-negative matrix in 𝑉 ∈ 𝑅^m×n^. Then 𝑉 could be decomposed into 2 non-negative matrices: 𝑊 ∈ 𝑅^m×k^ and 𝐻 ∈ 𝑅^k×n^, where 𝑘 represents the pre-selected factor number. Each element in 𝑊 matrix indicates the loading of a cell type on a corresponding factor, while elements of 𝐻 capture activity levels of these factor across tumor samples.

To assess the influence of the 𝑘 parameter on NMF outcomes, we ran the NMF analysis on the immune matrix using a range of values for 𝑘 (3 to 10), with 200 iterations for each value and a fixed seed. Following the guidelines suggested in the NMF vignette, we selected a factorization rank of 4 based on the following criteria: the first value where the cophenetic coefficient began to decrease after a high plateau, the rank exhibiting the highest dispersion value, and the rank at which the consensus silhouette score reached its peak. We considered each NMF factor to represent a distinct multicellular program. To identify the representative cell types associated with each program, we applied hierarchical clustering to the scaled 𝑊matrix.

To recover the NMF weights of these programs in validation samples, we used a custom NMF strategy that holds the pre-identified *W* matrix constant. This method initializes a new *H* matrix and iteratively updates it to find the best fit for the new data, minimizing the Euclidean distance between the new data matrix and the product of the fixed *W* and the new *H*. The resulting *H* matrix thus provides the weights levels of the four cellular programs for each sample.

### Assignment of scRNA-seq and bulk transcriptomic data to TIME subtypes

The relative abundance of immune cells of tumors for in-house validation set were calculated as aforementioned. For bulk transcriptomic data, the TME components were estimated by ssGSEA with representative gene sets. Then, we scaled the query data of the validation cohort to the original data and computed the Spearman’s correlation coefficient between the scaled proportions of each patient in the validation cohort and those in the original atlas. Each patient in the validation cohort was assigned to the group with the Spearman’s correlation coefficient above 0.25 with the corresponding group in the original atlas.

### Cell-cell interaction analysis

We applied CellChat v2^61^ to quantify the probability of signaling communication based on the expression of ligand-receptor pairs across cell types. CellChat employs the law of mass action to quantify communication probability and identify statistically significant interactions with permutation tests. To control for the false positivity, further filtered the interaction detected in at least 1/3 tumors in each group.

### Spatial transcriptomic analysis

The 10x Visium transcriptional data from NSCLC_Yan cohort was loaded as a scanpy^62^ object and pre-processed with 200 count per spot threshold. The probability of fine-grained cell subtype for each spot was estimated by cell2location^63^, a Bayesian model for spatial cell type mapping. To reduce the variation and ensure the accuracy, we used the matched scRNA-seq data from the same cohort as reference.

### Statistical analysis

All statistical analyses were performed in R software 4.4.0 and python 3.9. The statistical significance was set to p-value < 0.05.

## Data availability

All scRNA-seq and bulk transcriptomic datasets used in this study are accessible from their respective original publications, as detailed in **Table S1**. The Essen SKCM dataset has been deposited in the European Genome-phenome Archive under accession number EGAS50000000459.

## Code availability

The code for reproducing all the analyses in the study is available at https://github.com/zlin-bioinfo/PanCancer_ICI.

## Acknowledgments

This work was funded by the Israel Science Foundation (1543/21) and Azrieli Faculty Fellowship (to DA), by Deutsches Konsortium für Translationale Krebsforschung (ED03 to JCB) and Else-Kröner-Fresenius Stiftung (EFKS), University Medicine Essen Medical Scientist Academy (UMESciA) (to NS).

## Declaration of interests

JCB is receiving speaker’s bureau honoraria from Amgen, Pfizer, Recordati and Sanofi, is a paid consultant/advisory board member/DSMB member for Almirall, Boehringer Ingelheim, InProTher, ICON, MerckSerono, Pfizer, Regeneron, 4SC, and Sanofi. His group receives research grants from Merck Serono, HTG, IQVIA, and Alcedis. DA is a consultant to Link Cell Therapies. All other authors declare no competing financial interests.

## Author contributions

DA and JCB conceived the study and supervised the analyses. ZL performed all bioinformatics analyses. EP provided bioinformatics support. MC and NS performed the scRNA-seq. SU collected the samples from the patients. DA and ZL wrote the manuscript with inputs from JCB.

## Notes

### Competing Interest Statement

JCB is receiving speakers bureau honoraria from Amgen, Pfizer, Recordati and Sanofi, is a paid consultant/advisory board member/DSMB member for Almirall, Boehringer Ingelheim, InProTher, ICON, MerckSerono, Pfizer, Regeneron, 4SC, and Sanofi. His group receives research grants from Merck Serono, HTG, IQVIA, and Alcedis. DA is a consultant to Link Cell Therapies. All other authors declare no competing financial interests.

### Summary of Updates

Major revision with expanded dataset and revised analytical framework. This version includes: 1. Expanded cohort: 441 samples from 241 patients across ten cancer types (vs. 326 samples from 163 patients previously) 2. Revised deep-phenotyping framework: hierarchical reference-guided approach defining 77 immune and stromal subtypes (vs. integration-free framework with 83 cell states) 3. New central finding: identification of four conserved tumor immune microenvironment (TIME) subtypes with ~40% of tumors transitioning between states during therapy Addition of bulk transcriptomic validation cohort (1,988 tumors) Development of transition score for outcome prediction applicable to baseline tumors 4. Reframed focus on immunotype transitions as determinants of ICI response rather than cell subtype compositional changes 5. Updated figures and analyses throughout to reflect new framework and findings

