## Supplementary Figures for "Dynamic Tumor Immune Microenvironment Remodeling Predicts Response of Checkpoint Inhibitor Therapy"

### Supplementary Figure 1

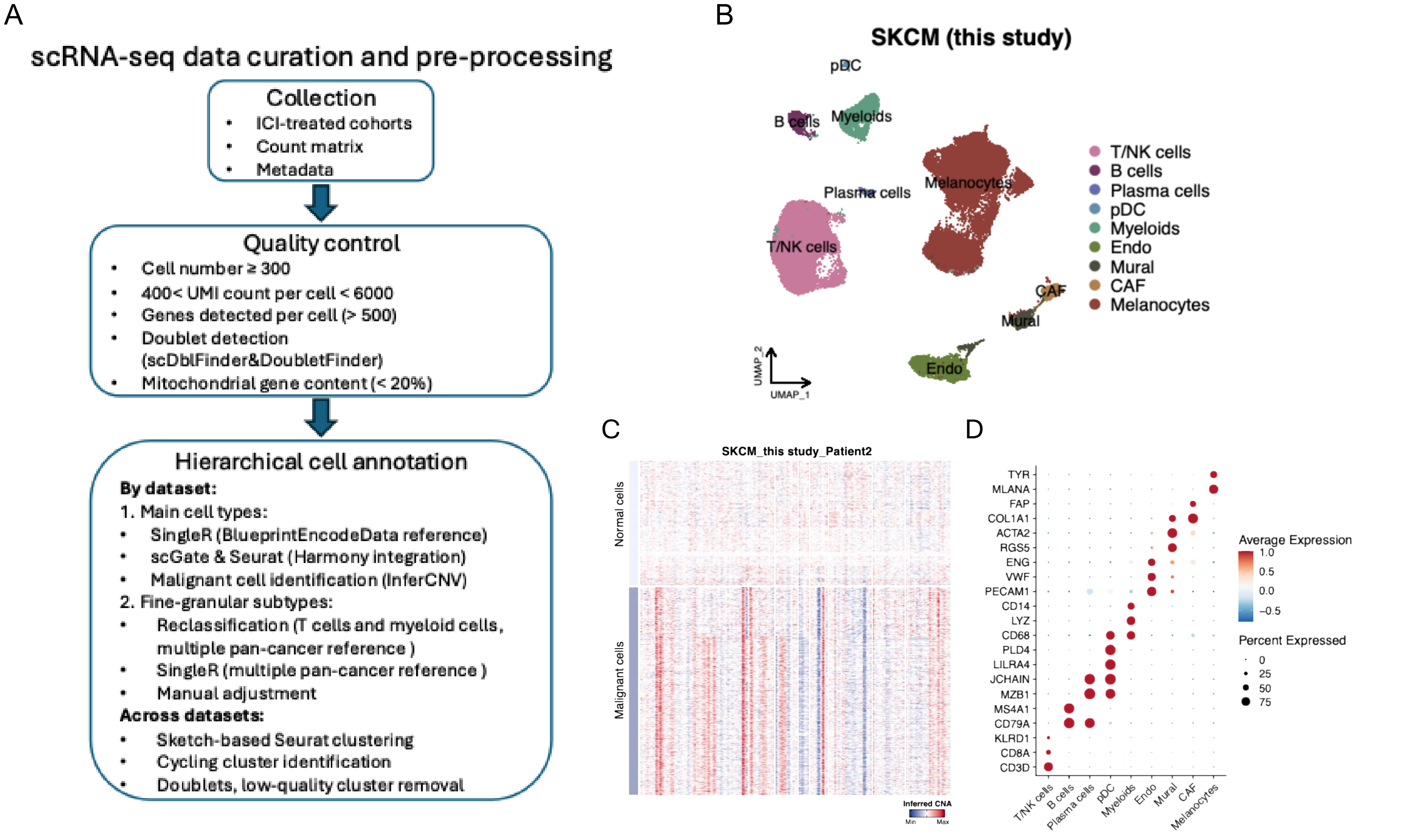


**Supplementary Figure 1. Overview of scRNA-seq data processing and quality control.**
**(A)** Schematic representation of data curation and hierarchical annotation workflow, including collection, quality control, and multilevel cell-type classification steps. **(B)** UMAP visualization of the in-house melanoma (SKCM) cohort, showing major immune, stromal, and malignant populations. (**C**) Inference of malignant cells by InferCNV, displaying copy-number aberrations (red, gain; blue, loss) for a representative patient, with malignant clusters annotated on the right. **(D)** Dot plot of canonical marker gene expression validating the classification of broad cell types in the in-house cohort.

### Supplementary Figure 2

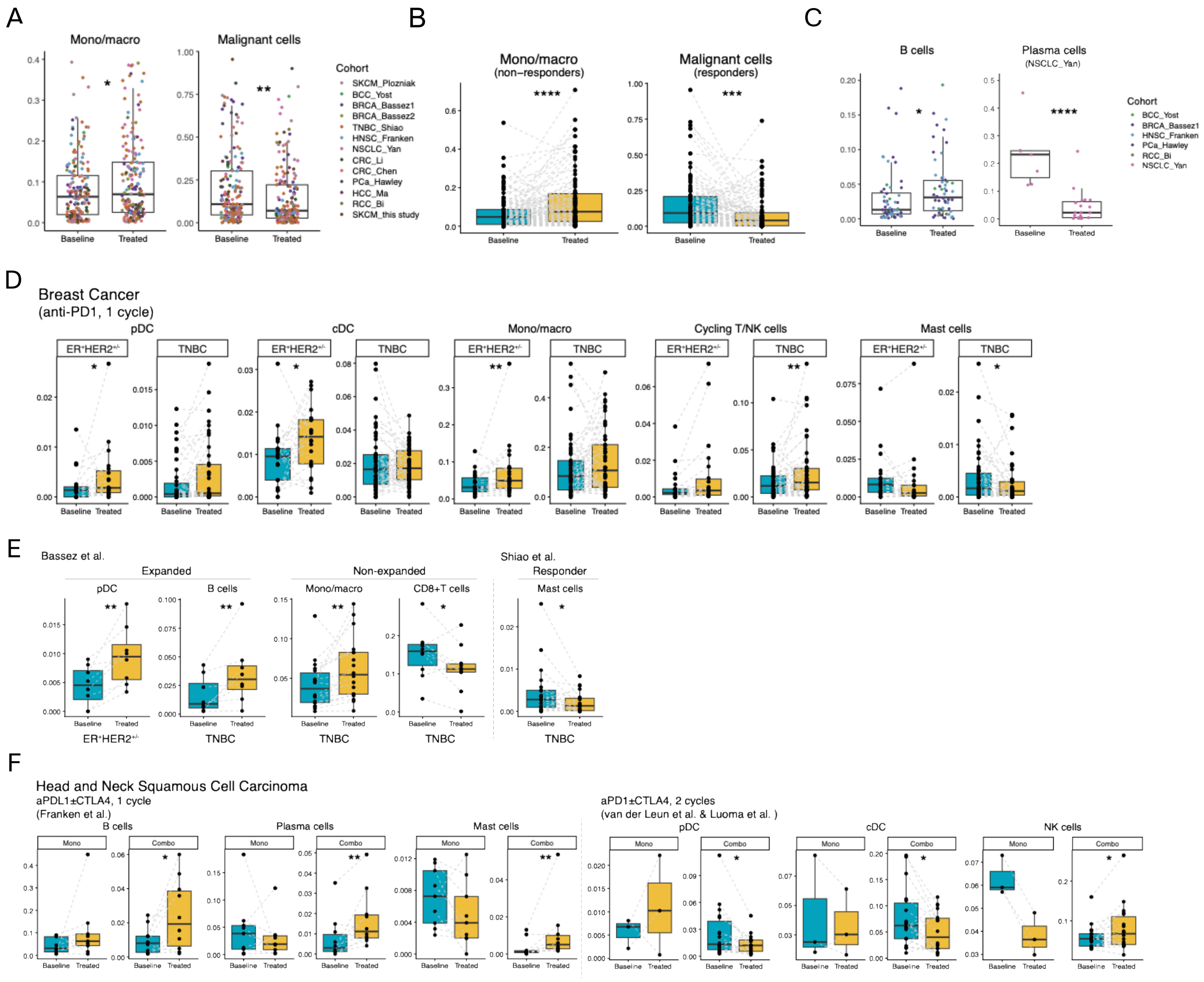


**Supplementary Figure 2. Cross-cohort analysis of immune-cell compositional changes during ICI therapy.**

**(A)** Comparison of monocyte/macrophage and malignant-cell abundance at baseline and on-treatment, with paired samples connected by dashed lines. **(B)** Compositional changes of monocytes/macrophages and malignant cells stratified by treatment response, based on standardized binary classifications. **(C)** Abundance of B cells and plasma cells at baseline and on-treatment across selected cohorts. **(D)** Subtype-specific immune shifts in breast cancer, showing changes in pDCs, cDCs, monocytes/macrophages, cycling T/NK cells, and mast cells after one anti-PD-1 treatment cycle. **(E)** Immune-composition changes linked to T-cell expansion status, highlighting cell-type differences between expanded and non-expanded groups. **(F)** Immune remodeling in head and neck squamous cell carcinoma (HNSCC) across treatment regimens and time points for anti-PD-1 ± anti-CTLA-4 therapy. Statistical comparisons were performed using paired Wilcoxon signed-rank tests ($p < 0.05$; *$p < 0.01$; **$p < 0.001$; ***$p < 0.0001$).

### Supplementary Figure 3

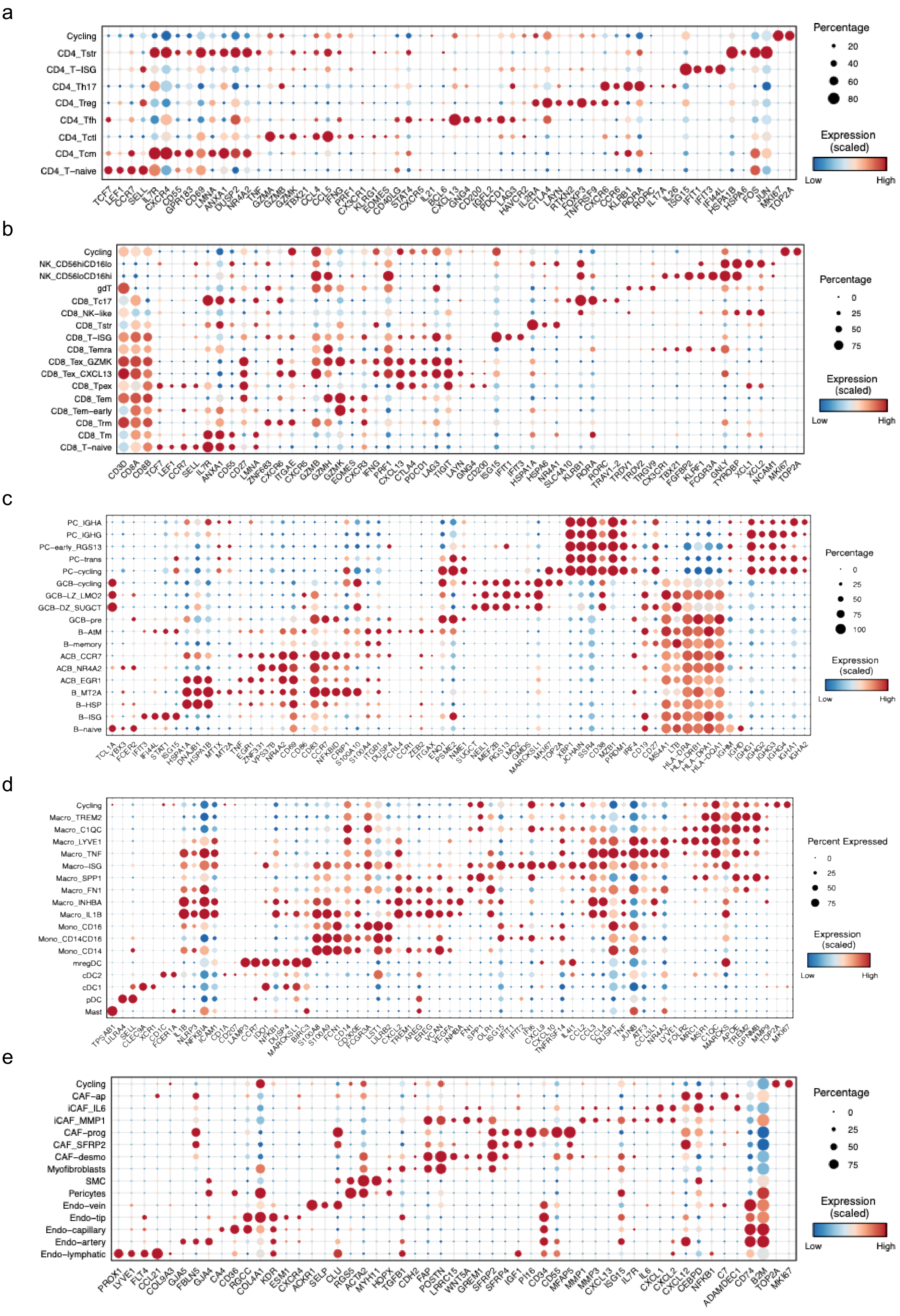


**Supplementary Figure 3. Validation of deep-phenotyping annotations.** Dot plots showing expression of canonical marker genes validating the classification of representative immune and stromal subtypes identified by the hierarchical annotation framework.

### Supplementary Figure 4

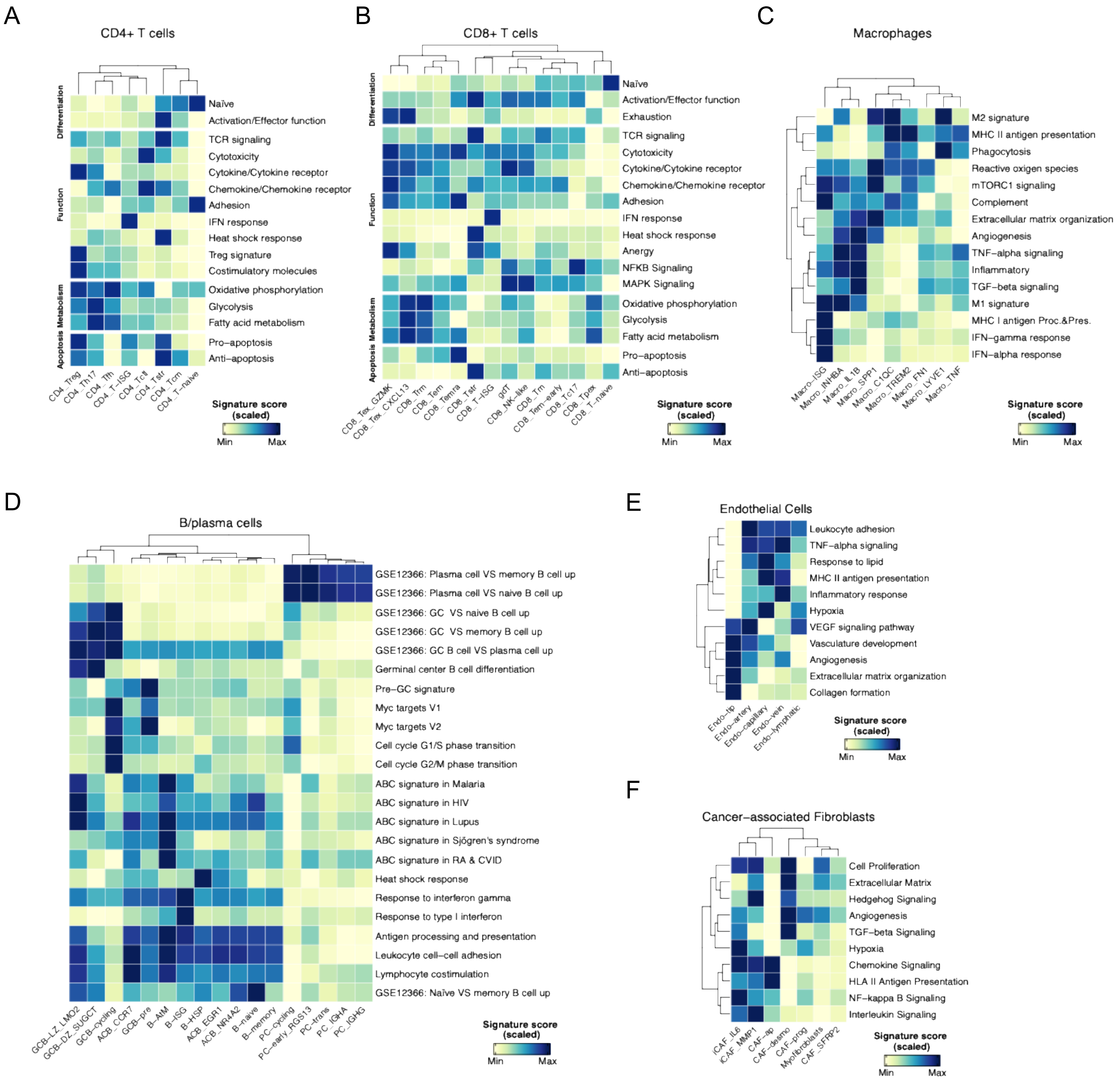


**Supplementary Figure 4. Functional validation of lineage-specific signatures.**
Heatmaps of curated functional signature scores confirming lineage-specific programs across major immune and stromal subtypes, including CD4$^{+}$ and CD8$^{+}$ T cells, B and plasma cells, monocytes/macrophages, endothelial cells, and cancer-associated fibroblasts (CAFs).

### Supplementary Figure 5

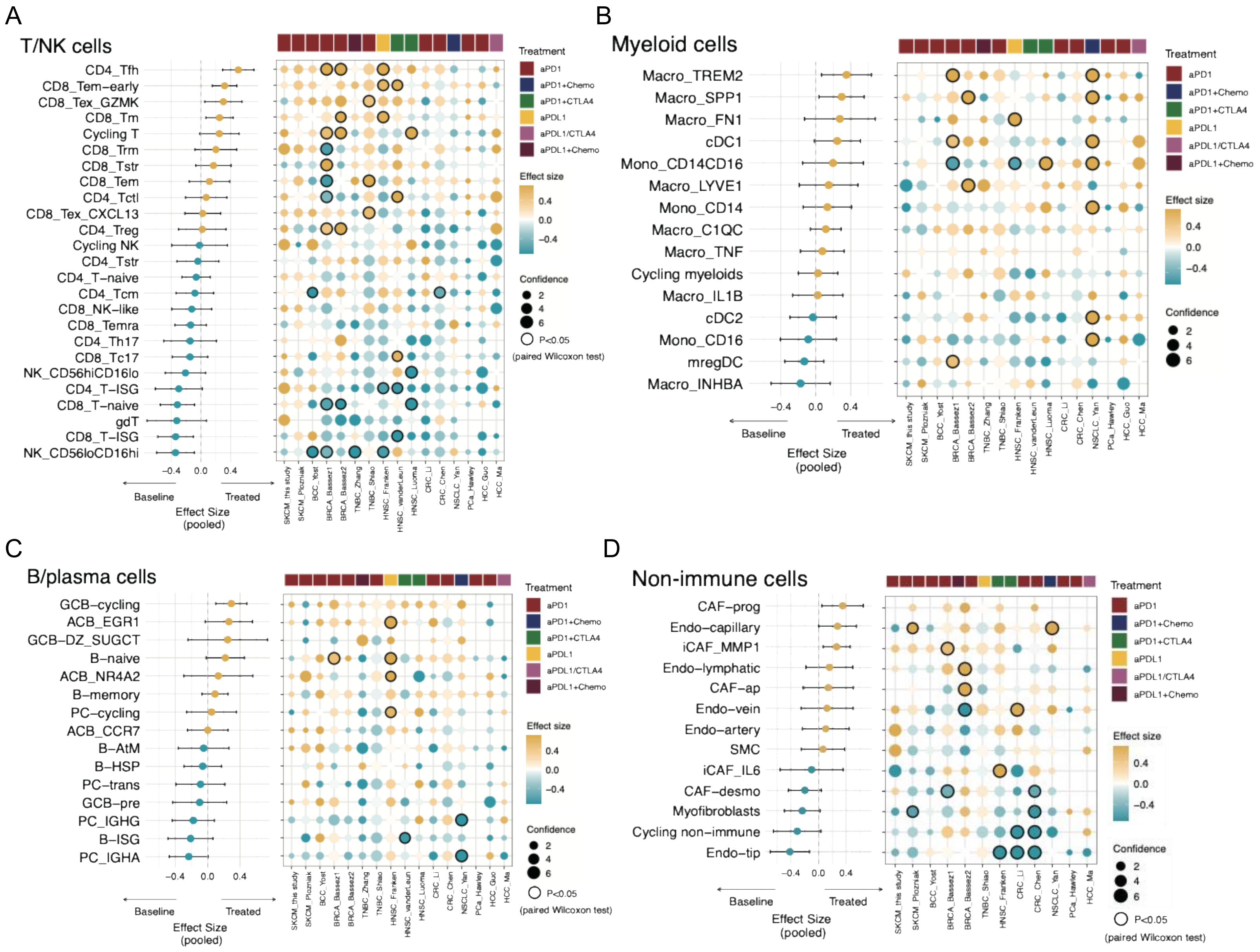


**Supplementary Figure 5. Pan-cancer compositional shifts following ICI therapy.**
Forest plots depicting pooled effect sizes (rank-biserial correlations) for changes in fine-grained immune and stromal subtypes between baseline and treated tumors across cancer types. Dot plots display study-specific abundance changes with confidence intervals. Paired Wilcoxon signed-rank tests were used for statistical evaluation.

### Supplementary Figure 6

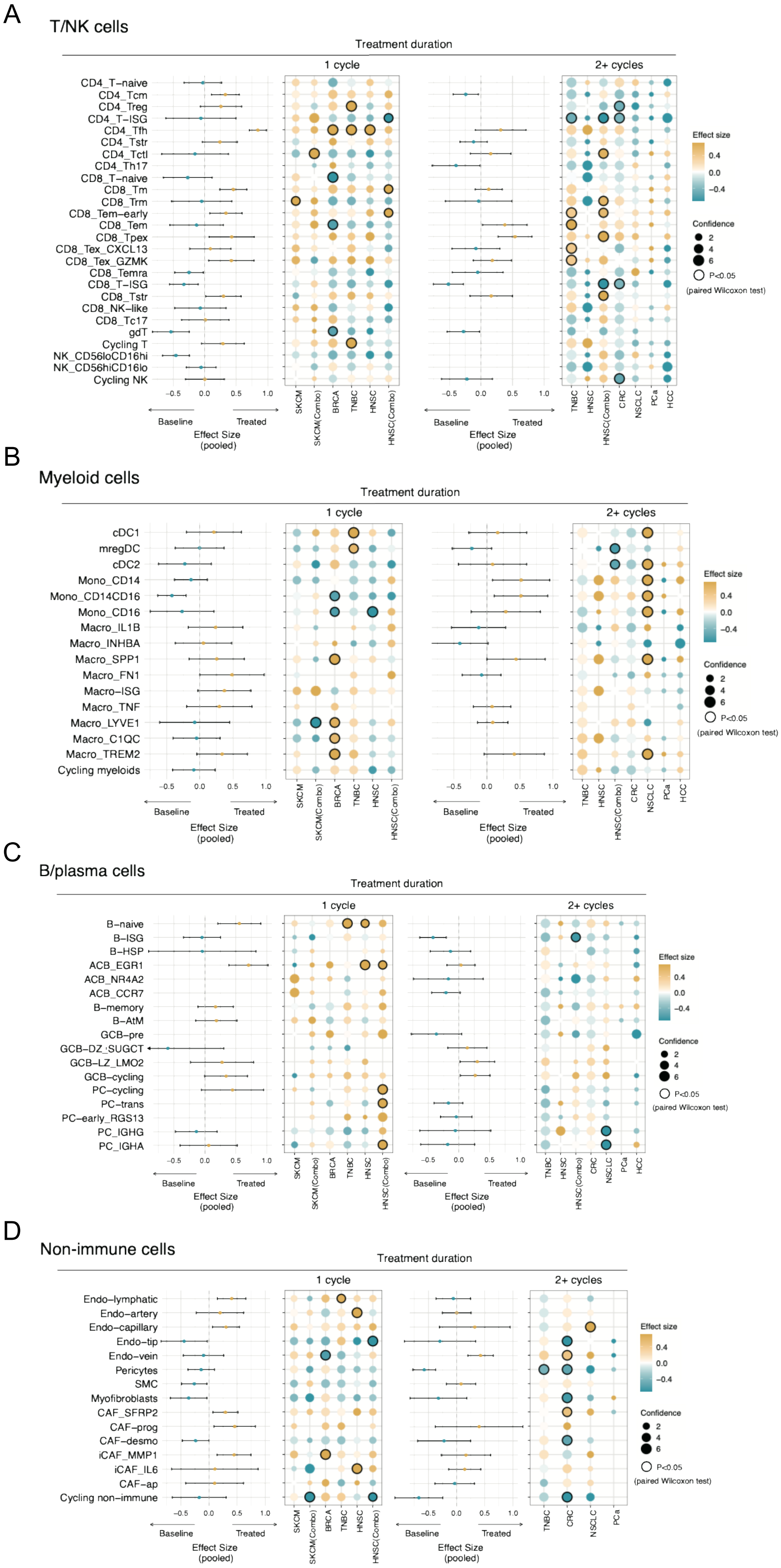


**Supplementary Figure 6. Temporal remodeling of the tumor immune microenvironment.** Forest plots summarizing compositional shifts in immune and stromal subtypes across cancer types, stratified by treatment duration (one versus ≥2 cycles). Dot plots indicate cohort-specific abundance changes and 95% confidence intervals. Paired Wilcoxon signed-rank tests were used for statistical assessment.

### Supplementary Figure 7

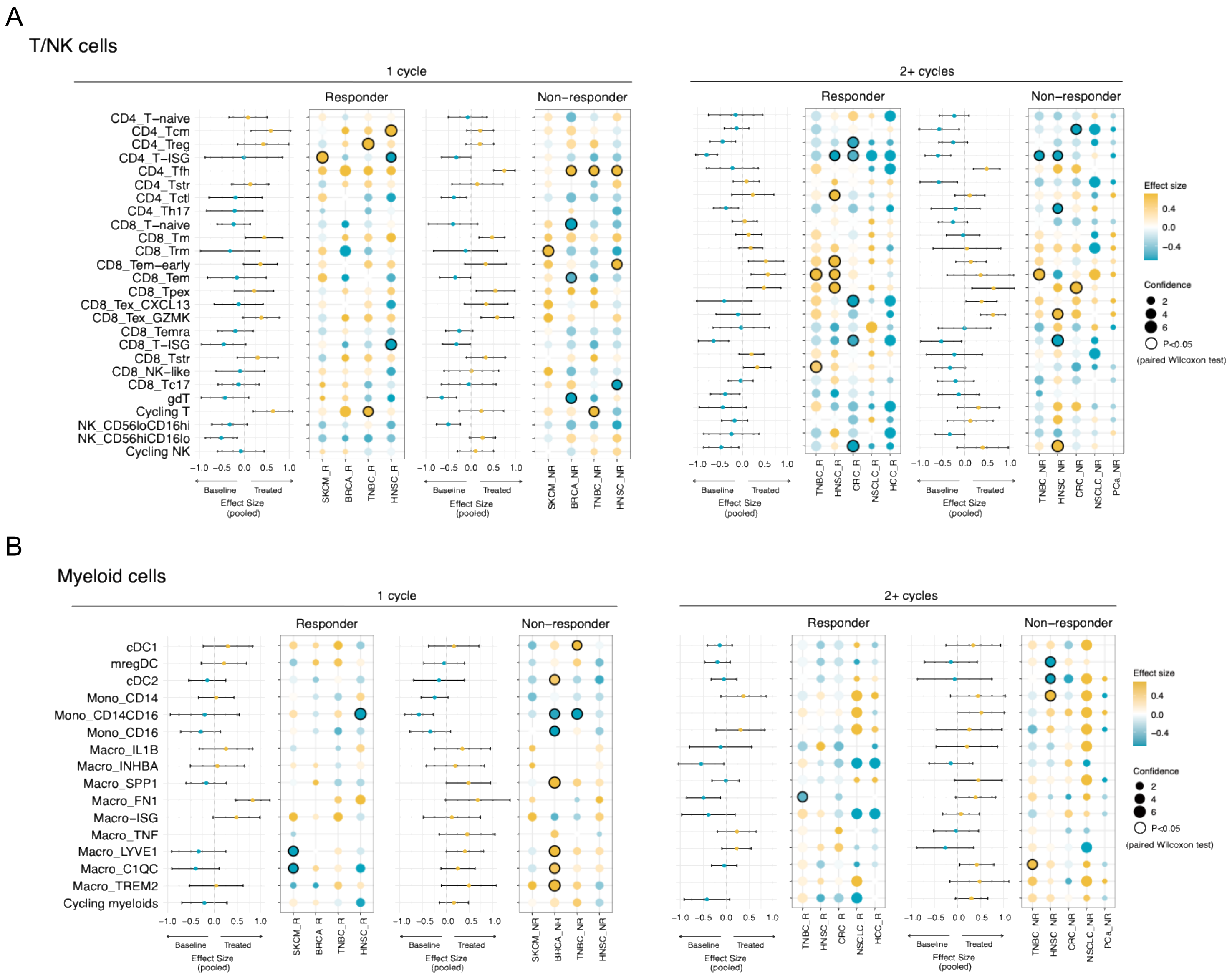


**Supplementary Figure 7. Association of T/NK and myeloid remodeling with treatment response.** Forest plots showing pooled effect sizes and study-specific abundance changes of T/NK and myeloid subtypes across cancer types, stratified by treatment duration and clinical response. Statistical significance was determined by paired Wilcoxon signed-rank tests.

### Supplementary Figure 8

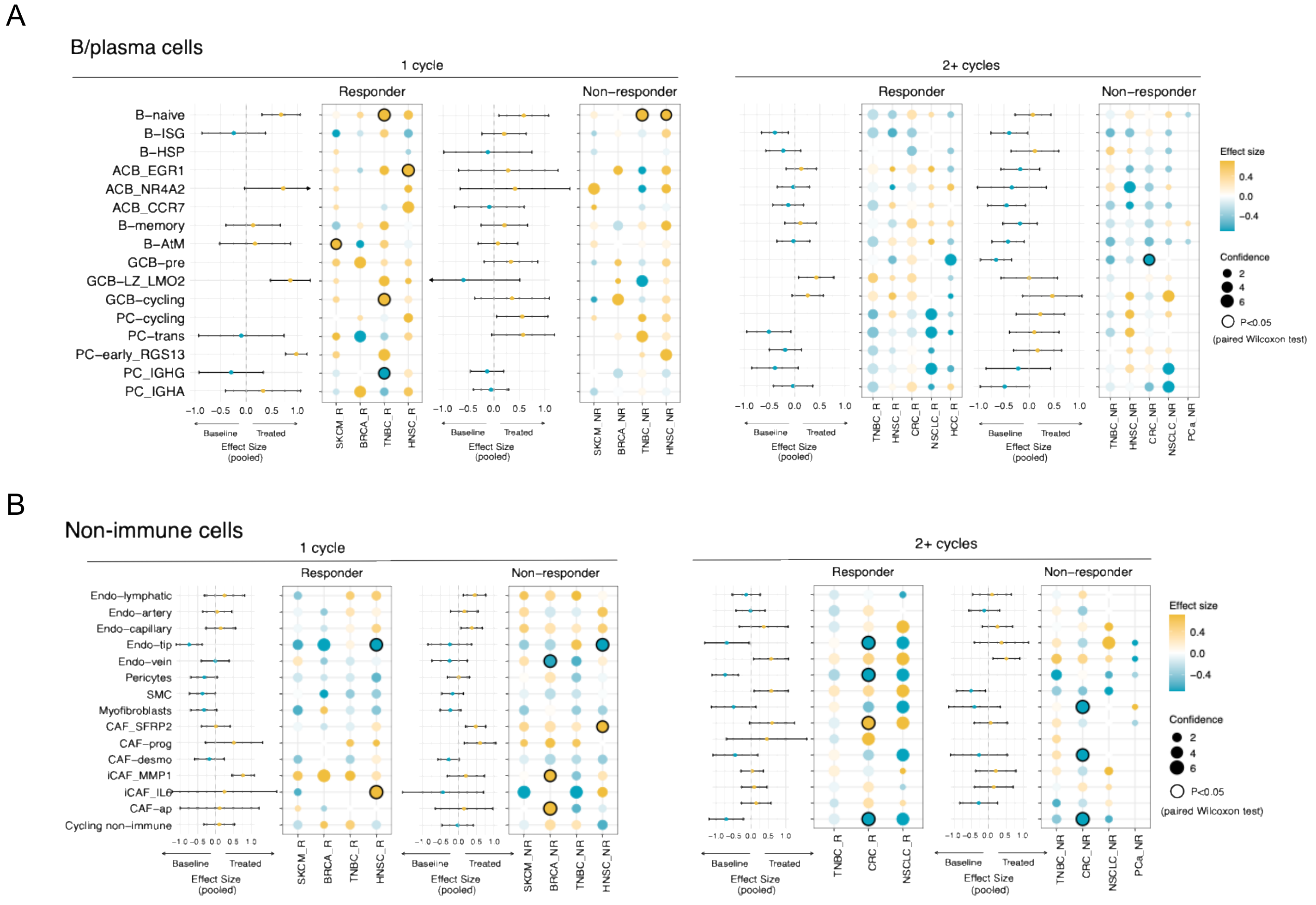


**Supplementary Figure 8. Response-associated remodeling of B, plasma, and non-immune compartments.** Forest plots showing pooled and study-specific abundance changes in B/plasma cell and non-immune stromal subtypes (endothelial and fibroblast lineages) across cancer types, stratified by treatment duration and response. Statistical testing by paired Wilcoxon signed-rank tests.

### Supplementary Figure 9

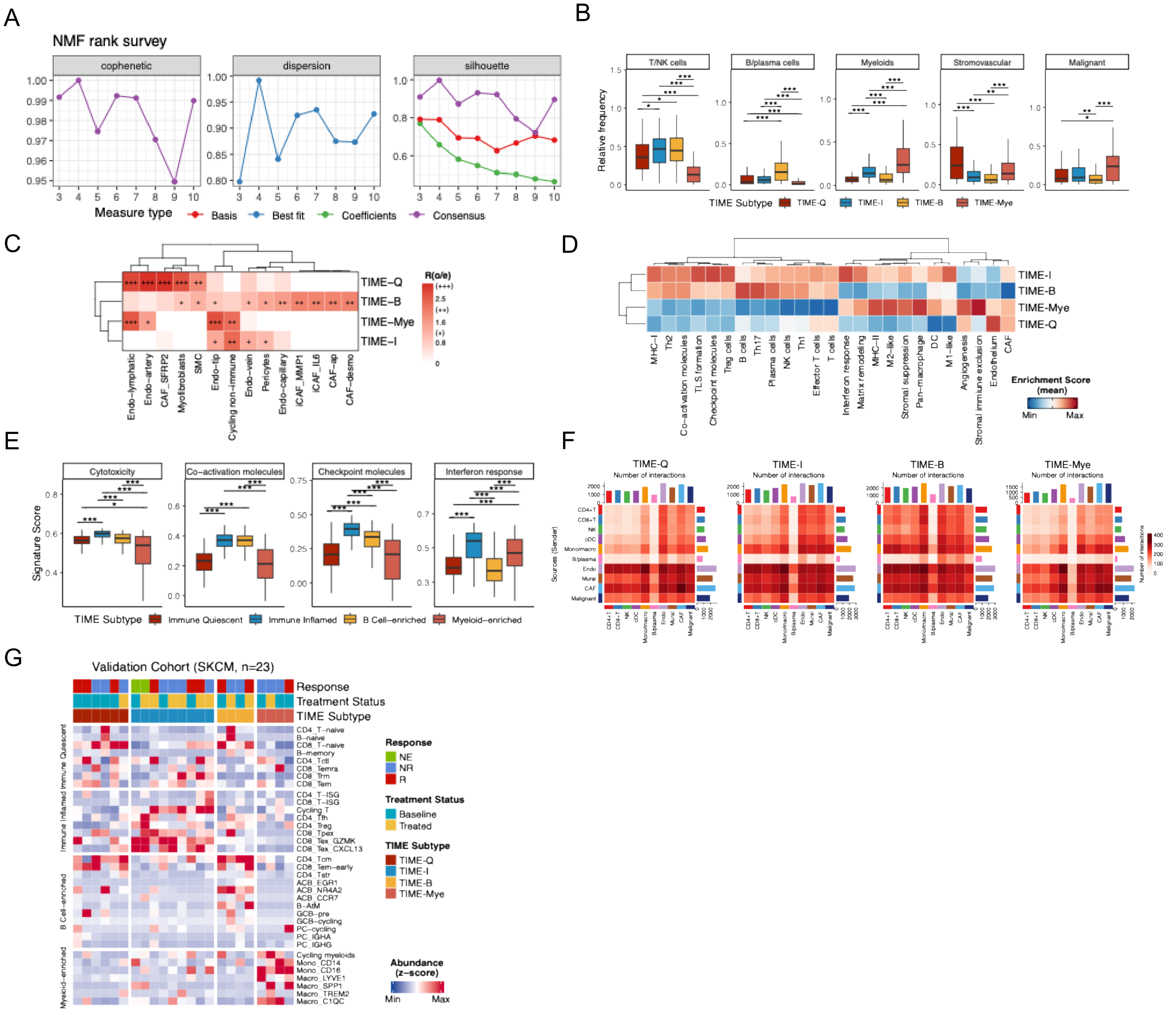


**Supplementary Figure 9. Characterization and validation of TIME subtypes in bulk transcriptomic cohorts. (A)** NMF rank survey metrics used to select optimal factorization rank. **(B)** Boxplots comparing the proportions of broad cell lineages across TIME subtypes. Statistical testing by Wilcoxon signed-rank test. **(C)** Heatmap showing distribution preferences of non-immune cell subtypes across TIME subtypes. **(D)** Average pathway activity scores across TIME subtypes. **(E)** Boxplots comparing functional signature scores between TIME subtypes. Statistical testing as in (B). **(F)** Heatmap comparing TIME subtypes with carcinoma ecotypes (Thorsson et al.). **(G)** Heatmaps showing predicted cell–cell interaction strength (CellChat) for each TIME subtype. **(H)** Heatmap showing relative abundance of representative immune populations in the in-house melanoma cohort, validating TIME classification. *p < 0.05, **p < 0.01, ***p < 0.001, ****p ≤ 0.0001

### Supplementary Figure 10

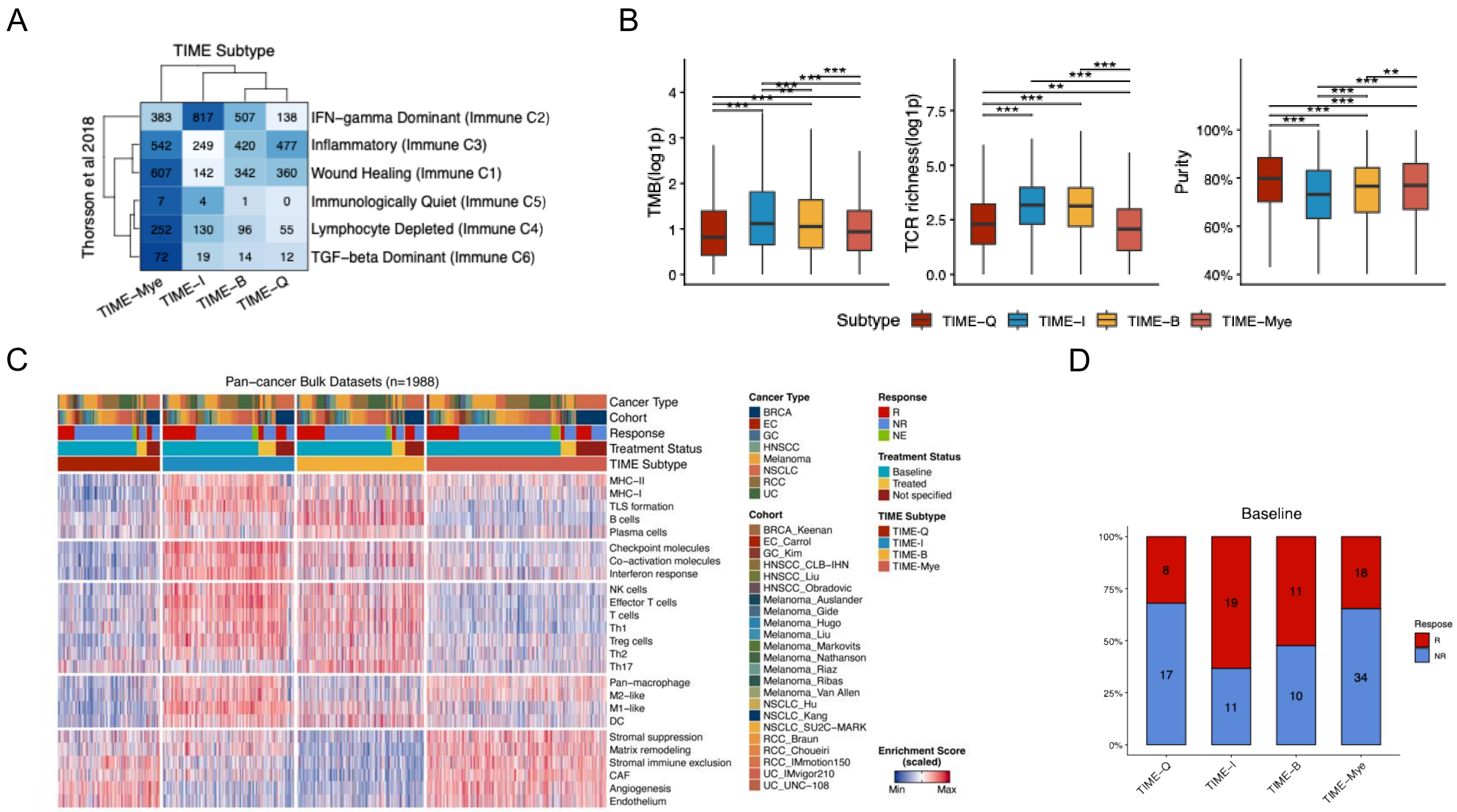


**Supplementary Figure 10. Transfer of TIME subtypes to ICI-treated bulk transcriptomic datasets. (A)** Heatmap comparing TIME subtypes with carcinoma ecotypes (Thorsson et al.). **(B)** Boxplots comparing tumor mutational burden (TMB), T-cell receptor (TCR) richness, and tumor purity across TIME subtypes. Statistical testing by Wilcoxon signed-rank test. **(C)** Heatmap of 1,988 ICI-treated tumors classified into four distinct TIME subtypes. **(D)** Bar plot showing baseline response rates across TIME subtypes. *p < 0.05, **p < 0.01, ***p < 0.001, ****p ≤ 0.0001

### Supplementary Figure 11

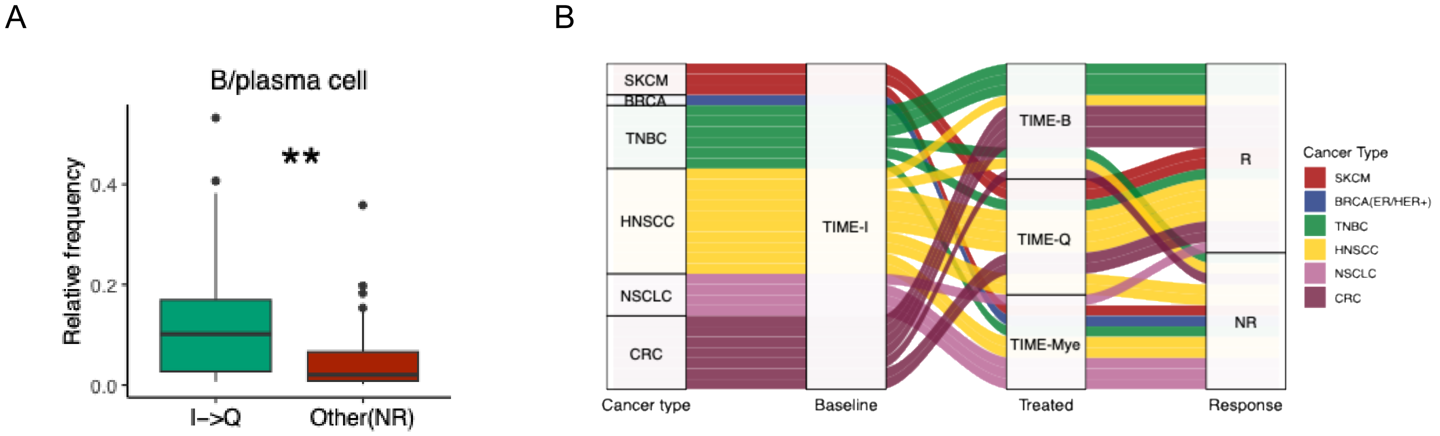


**Supplementary Figure 11. Cancer type–specific TIME-I transition patterns and B-cell enrichment in responsive tumors. (A)** Alluvial diagram illustrating TIME subtype transitions originating from baseline TIME-I tumors across multiple cancer types following ICI treatment. Each flow represents a paired pre- and post-treatment sample, with color indicating the destination subtype. **(B)** Boxplots comparing B-cell and plasma-cell abundance between responsive TIME-Q tumors that transitioned from TIME-I and non-responsive TIME-Q tumors. Two-tailed unpaired Wilcoxon tests.

### Supplementary Figure 12

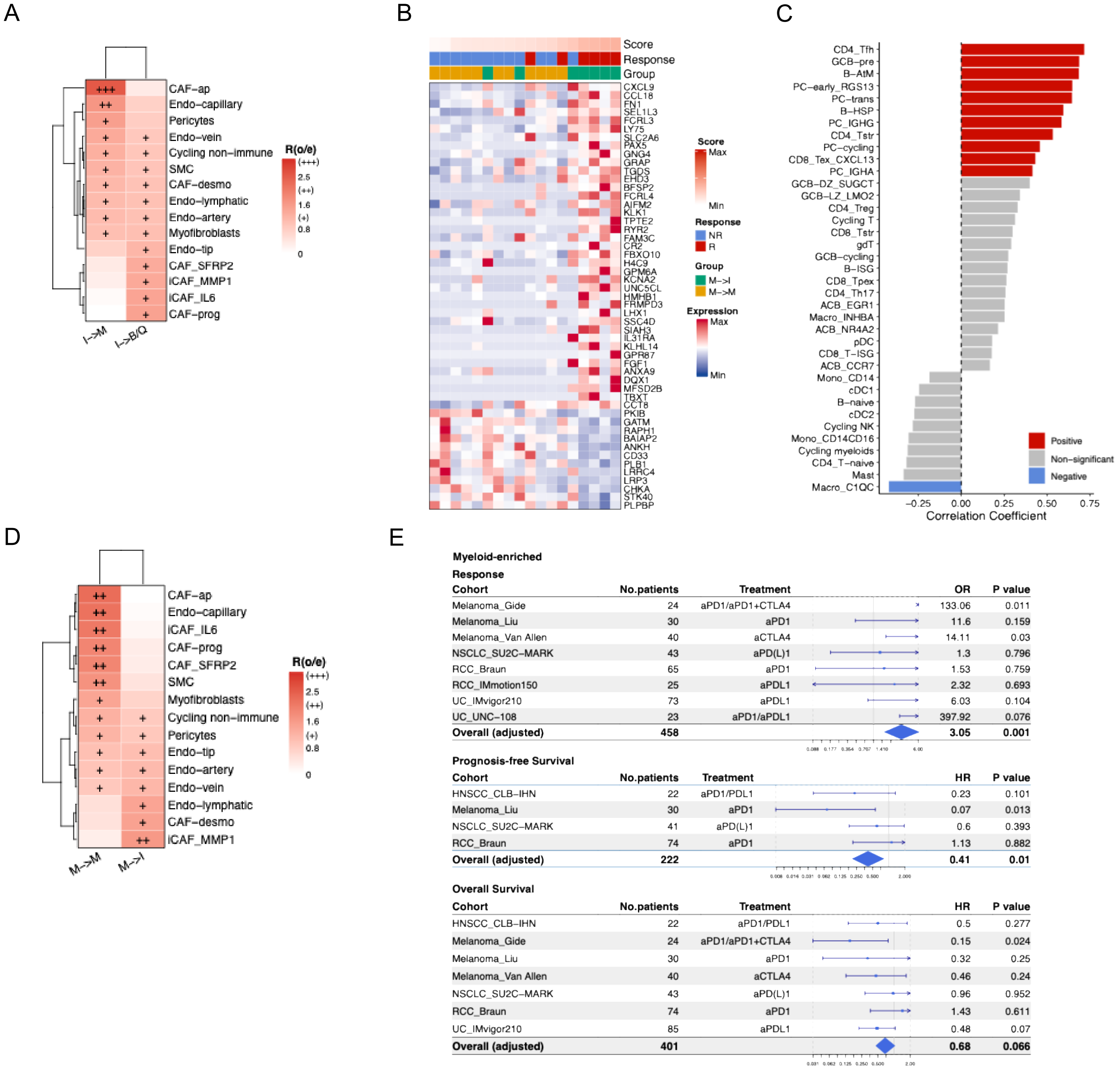


**Supplementary Figure 12. Baseline determinants of myeloid transitions and therapeutic outcomes.**
**(A)** Heatmap showing the distribution of non-immune stromal and vascular cell types in baseline TIME-I tumors stratified by transition fate. **(B)** Heatmap of the top 50 upregulated and 50 downregulated genes distinguishing baseline TIME-Mye tumors that transitioned toward TIME-I versus those remaining myeloid-stable. **(C)** Forest plots showing predictive performance of the TIME-Mye transition score for treatment response, PFS, and OS in baseline bulk ICI transcriptomic cohorts. **(D)** Bar plot showing Spearman correlations between TIME-Mye transition scores and immune-cell subtype abundance in baseline tumors. **(E)** Heatmap of non-immune cell-type distribution in baseline TIME-Mye tumors stratified by transition fate.
